# Single-cell transcriptomic profiling of progenitors of the oligodendrocyte lineage reveals transcriptional convergence during development

**DOI:** 10.1101/186445

**Authors:** Sueli Marques, Darya Vanichkina, David van Bruggen, Elisa M. Floriddia, Hermany Munguba, Leif Väremo, Stefania Giacomello, Ana Mendanha Falcão, Mandy Meijer, S Samudyata, Simone Codeluppi, Åsa K. Björklund, Sten Linnarsson, Jens Hjerling-Leffler, Ryan J. Taft, Gonçalo Castelo-Branco

## Abstract

*Pdgfra+* oligodendrocyte precursor cells (OPCs) arise in distinct specification waves during embryogenesis in the central nervous system (CNS). It is unclear whether there is a correlation between these waves and different transcriptional oligodendrocyte (OL) states at adult stages. Here we present a bulk and single-cell transcriptomics resource providing insights on how transitions between these states occur. We show that E13.5 *Pdgfra+* populations are not OPCs, exhibiting instead hallmarks of neural progenitors. A subset of these progenitors, which we refer as pre-OPCs, rewires their transcriptional landscape, converging into indistinguishable OPC states at E17.5 and post-natal stages. P7 brain and spinal cord OPCs present similar transcriptional profiles at the single-cell level, indicating that OPC states are not region-specific. Postnatal OPC progeny of E13.5 *Pdgfra+* have electrophysiological and transcriptional profiles similar to OPCs derived from subsequent specification waves. In addition, lineage tracing indicates that a subset of E13.5 *Pdgfra+* cells also originate cells of the pericyte lineage. In summary, our results indicate that embryonic *Pdgfra+* cells are diverse and give rise at post-natal stages to distinct cell lineages, including OPCs with convergent transcriptional profiles in different CNS regions.

## Introduction

Oligodendrocytes (OLs) are one of the most abundant cell types in the CNS. OLs have been classically described as support cells for neurons, responsible for the insulation of axons and enabling rapid saltatory conduction, although recent findings suggest their involvement in other processes (Micu, et al., 2016; Nave and Werner, 2014). OLs arise from the differentiation of oligodendrocyte precursor cells (OPCs). Since OLs and OPCs are found throughout the CNS, they were originally thought to be derived from embryonic neural progenitors from nearby ventricular zones or from radial glial cells in all CNS regions (Hirano and Goldman, 1988). However, *in vitro* experiments with cells/explants suggested a restricted ventral origin of OPCs in the embryonic rat spinal cord (Warf, et al., 1991). OPCs expressing platelet-derived growth factor receptor alpha (*Pdgfra*) appeared to arise exclusively from ventral domains of the CNS, followed by migration to dorsal regions (Pringle and Richardson, 1993). This led to the hypothesis of a single embryonic lineage for OPCs, arising at embryonic day (E)12.5 from progenitor domains dependent on sonic hedgehog (Shh) (Richardson, et al., 2000). Nevertheless, cells expressing *Plp/dm-20* were also observed to give rise to OLs (Spassky, et al., 2000), suggesting that other progenitors in the dorsal/ventral axis of the CNS could be alternative sources of OPCs. Indeed, knockdown of transcription factors involved in the specification of ventral spinal cord domains and lineage tracing studies uncovered a subset of OPCs originating from the mouse dorsal spinal cord at E14.5-E15.5 (Cai, et al., 2005; Fogarty, et al., 2005; Vallstedt, et al., 2005). Dorsal-derived spinal cord OPCs were not dependent on Shh for their specification (Cai, et al., 2005; Vallstedt, et al., 2005). Moreover, they populated specific regions of the spinal cord, while OPCs from the ventral-derived neuroepithelium gave rise to the vast majority of OLs throughout the adult spinal cord (Tripathi, et al., 2011; Fogarty, et al., 2005). A similar pattern of OL specification was found to operate in the mouse forebrain. The first OPCs were shown to be formed at E12.5 from *Nkx2.1* expressing precursors in the medial ganglionic eminence, followed by a subsequent wave from precursors expressing *Gsx2* in the lateral and medial ganglionic eminences at E15.5 (Kessaris, et al., 2006). The last wave of cortically derived *Emx1*-expressing precursors begins developing at birth and eventually becomes the major contributor to the postnatal OPC pool (Kessaris, et al., 2006).

It is unclear whether diverse populations and developmental waves of OPC in the CNS generate identical or distinct OLs. Richardson and colleagues specifically ablated OLs generated from three brain developmental waves in mice, with no gross behaviour abnormalities observed, most likely due to functional compensation of the lost populations by OLs from non-ablated regions (Kessaris, et al., 2006). However, adult OPCs have been subdivided into different sub-populations with diverse electrophysiological (Clarke, et al., 2012; Tripathi, et al., 2011; Karadottir, et al., 2008), cycling (Jang, et al., 2013; Young, et al., 2013; Clarke, et al., 2012), myelinating (Vigano, et al., 2013) and remyelination (Crawford, et al., 2016) properties. In addition, spinal cord postnatal OPCs give rise to OLs that present a higher myelin sheet length than OLs from the cortex (Bechler, et al., 2015). By performing single-cell RNA sequencing on ∼5000 cells of the OL lineage in juvenile and adult mice, we have also shown that there are at least six distinct mature OL cell states (Marques, et al., 2016), suggesting possible functional heterogeneity. These recent findings challenge a putative homogeneity of OPCs, highlighting the need to reexamine whether the embryonic waves of OPC specification give rise to redundant OLs or are assigned to specific mature OL states/populations.

In order to investigate whether OPCs are transcriptionally heterogeneous, we analyzed the transcriptome of mouse forebrain and spinal cord *Pdgfra+* cells at E13.5, E17.5, P7 and juvenile/adult stages, by bulk and single-cell RNA-Seq. E13.5 *Pdgfra+* cells were found to constitute not one but six distinct cell states within the OL (which we refer as pre-OPCs), pericyte and possibly other lineages. Interestingly, E13.5 *Pdgfra+* did not express “hallmark” genes of OPCs, but rather of neural progenitors. Embryonic OL progenitors expressed several patterning transcription factors (TFs), whose expression was greatly attenuated at E17.5 and post-natal stages, when a common transcriptional profile compatible with electrophysiological capacity emerged. Single-cell RNA-Seq analysis also indicated that spinal cord and brain OPCs had undistinguishable profiles after birth. Post-natal OPCs in the postnatal forebrain derived from distinct embryonic specification waves had similar electrophysiological and single-cell transcriptomic profiles, highlighting spatial and temporal transcriptional convergence in the transition between embryonic pre-OPCs and OPCs during development.

## Results

### Transcriptional profiles of *Pdgfra+* cells during CNS development

We performed stranded total RNA-Seq of cells expressing *Pdgfra*, a widely used marker for OPCs in CNS, isolated by FACS sorting GFP+ populations from the forebrain or spinal cord of *Pdgfra*-H2B-GFP mice (Klinghoffer, et al., 2002) at E13.5 and P7 (Figure 1a and Figure S1). Differential gene expression analysis revealed increased levels of differentiation/myelination related genes (*Mbp, Plp1, Mag* and *Mog*, among others) in *Pdgfra*+/GFP cells at P7 relative to E13.5, and in the spinal cord when compared to the brain (*Opalin* and *Plp1*, Figure 1b; Table S1). P7 spinal cord *Pdgfra*+/GFP cells were characterised by higher expression of genes corresponding to later stage differentiation (*Mog, Mal, Mag*) when compared to P7 brain, which in contrast exhibit increased expression of *Pdgfra, Sox2 and Sox9* (Figure 1c and Table S1). This pattern of expression would suggest that OPCs are more prone to differentiation in the spinal cord, consistent with myelination occurring earlier in the spinal cord compared to brain (Marques, et al., 2016; Coffey and McDermott, 1997).

**Fig. 1.**
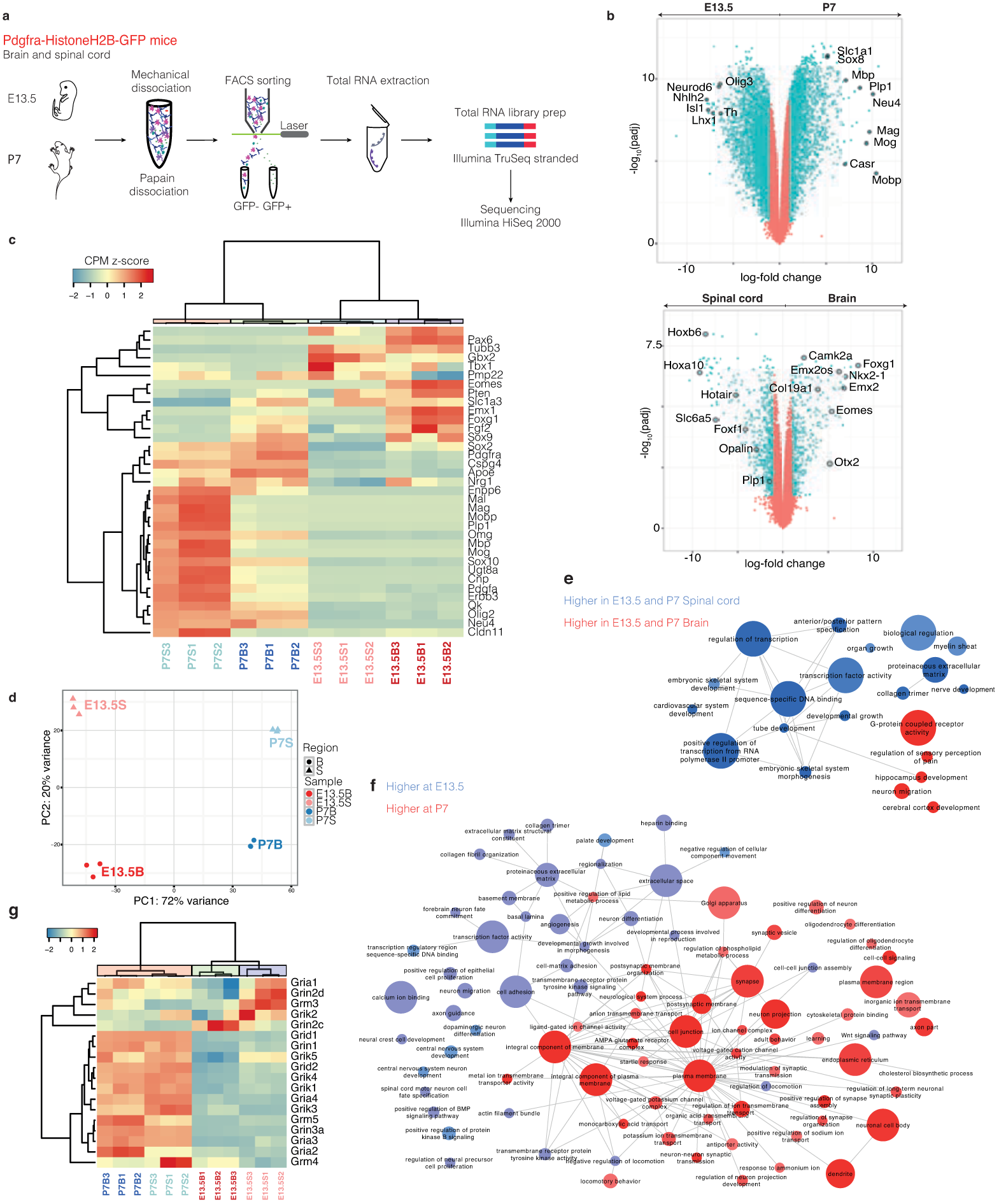
Temporal and spatial transcriptional heterogeneity of Pdgfra+/GFP cells.

a) Schematic of OPC purification for bulk RNA sequencing;
b) Volcano plots of Gencode-annotated genes depicting differential expression between forebrain vs spinal cord, and E13.5 versus P7;
c) Heatmap showing hierarchical clustering of bulk samples based on normalised gene expression (counts per million mapped reads, cpmm) of selected genes involved in oligodendrocyte lineage progression;
d) Principal component analysis of bulk RNA-Seq of Pdgfra+/GFP cells from E13.5 and P7 forebrain and spinal cord;
e/f) Gene ontology analysis of enriched biological functions overrepresented in either forebrain vs spinal cord (e), or E13.5 vs. P7 (f);
g) Hierarchical clustering of bulk samples based on normalised gene expression (cpmm) of a subset of glutamate receptors expressed in Pdgfra+/GFP cells.

Principal component analysis confirmed that *Pdgfra+/*GFP populations were more distinct temporally than spatially (Figure 1d). Gene ontology (GO) analysis (Table S1) indicated that, apart from genes involved in differentiation/myelination, spinal cord cells were enriched in genes with transcription factor activity (Figure 1e and Figure S2), consistent with the expression of patterning transcription factors involved in anterior/posterior regionalization (Figure 1b). Transcription factor activity was also enriched in E13.5 *Pdgfra*+/GFP cells compared to P7 (Figure 1f). Post-natal *Pdgfra*+ZGFP cells, in particular in the forebrain, were enriched in genes involved in networks regulating ion channel complex and transport, and synaptic transmission, among others (Figure 1f and Figure S2a). Some of the contributions to these postnatal signatures were potassium and sodium ion channels, as well as glutamate receptor subunits and GABA receptors (Figure 1g and Figure S2b). In contrast to post-natal cells, embryonic *Pdgfra*+/GFP cells were enriched in genes involved in a myriad of unrelated processes, such as neural precursor and neuron development/specification, extracellular matrix and collagen organization, basal lamina and angiogenesis (Figure 1f and Figure S2c). This diversity of biological function was intriguing, since these cells are thought to give rise exclusively to cells of the OL lineage.

### Transcriptional networks operating in embryonic and post-natal *Pdgfra+* cells

Many transcription factors involved in patterning and cell specification commitment were expressed in *Pdgfra*/GFP+ cells at E13.5 and substantially downregulated or absent at P7 (Figure 2a and Table S1). In contrast, transcription factors that have been shown to have roles in OL differentiation only emerged at P7. In order to further investigate the transcriptional transitions operating in *Pdgfra*/GFP+ cells, we performed upstream regulator analysis using QIAGEN’s Ingenuity Pathway Analysis (IPA®, www.qiagen.com/ingenuity) (Figure 2b,c; Figure S3 and Table S2), considering only experimentally validated interactions. Spinal cord cells presented an intricate network of *Hox* transcription factors, as well as *Pax8*, *Foxf1* and *Lbx1*. IPA analysis predicted that this network is controlled by the activity of the transcription factor *Hoxa10*, but also of two epigenetic modulators, *Kat6a* (Myst3/ Moz) and *Kmt2a* (Mll1). Brain *Pdgfra+/*GFP cells presented in contrast a network of anterior transcription factors, such as *Emx1/2, Otx1/2, Dlx1*/*2, Lhx2, Nkx2.1*, *Eomes* among others (Figure 2b and Figure S3). *Sox2* and *Pax6* were enriched in the brain and predicted to regulate some of these transcriptional networks, along with epigenetic modulators belonging to the family of Polycomb proteins, such *Bmi1* (*Commd3-Bmi1*) and *Phc2* (Figure S3). This suggests that these chromatin regulators, which have not been previously implicated in OL lineage progression, might primarily act in these cell populations to regulate expression of patterning regulatory networks.

**Fig. 2.**
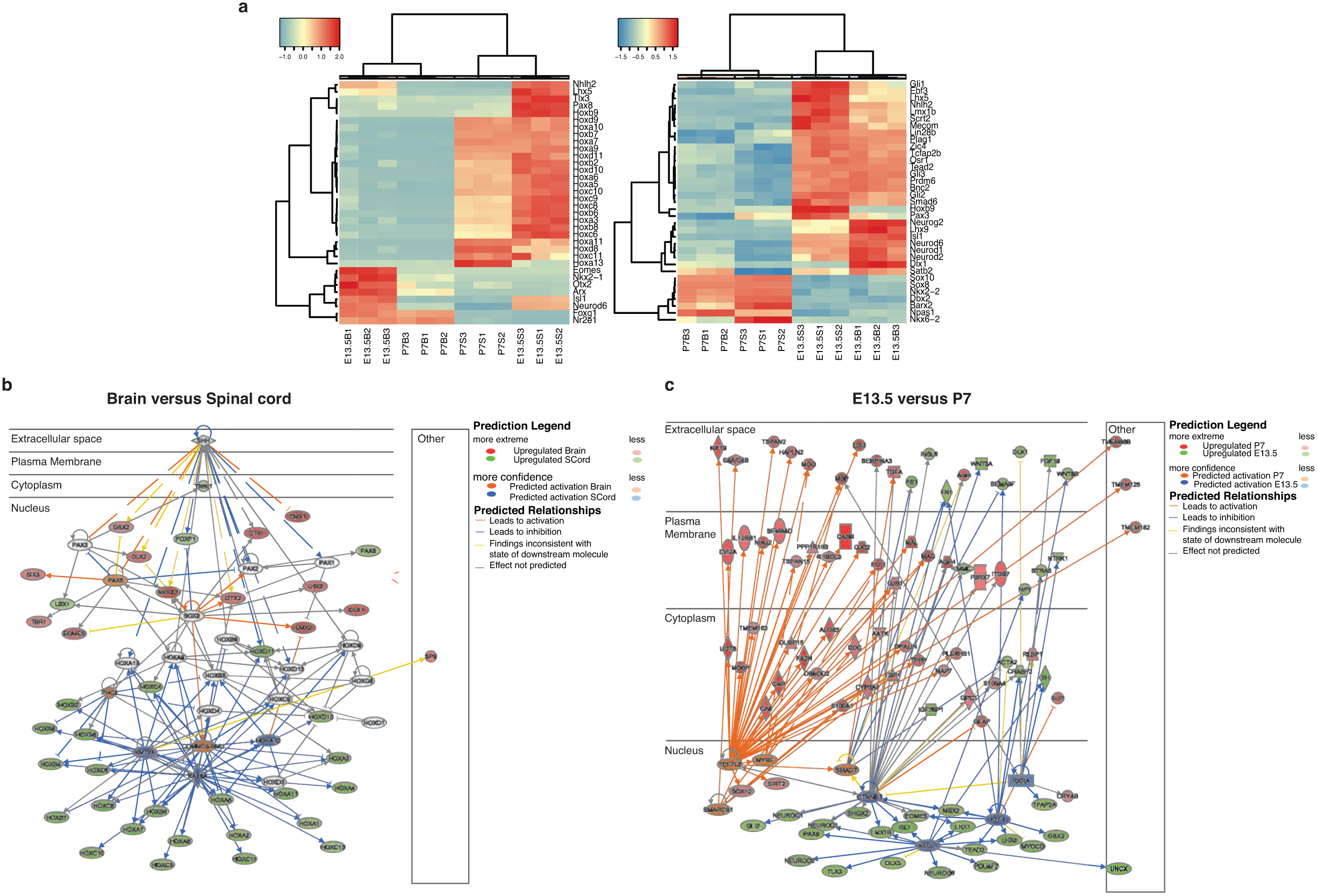
Transcriptional networks operating in embryonic and post-natal Pdgfra+ cells.

Hierarchical clustering of bulk samples based on normalised gene expression (cpmm) of transcription factors annotated in animalTFDB (Zhang et al., 2015); b-e) Network of genes controlled by the IPA- identified upstream regulators: (b) Commd1-Bmi1, Kat6a, Kmt2a, Phc2, Shh, Sox2, and Hox and Pax family members in the bulk brain and spinal cord RNA-seq datasets; (c) Ascl1, Cttnb1, Klf4, Myrf, Rxra, Smad7, Smarcb1, and Tcf7l2 in the bulk E13.5 and P7 RNA-seq datasets.

The transcription factor *Klf4* and the nuclear receptor *Rxra* were enriched and predicted to be active at E13.5 (Figure 2c and Figure S3). Other transcription factors such as *Ascl1* (Sugimori, et al., 2008) and beta-catenin (*Ctnnb1*) were also predicted to be activators of this network at E13.5. However, *Ctnnb1* was not differentially expressed between E13.5 and P7, while *Ascl1* was paradoxically enriched at P7 (Table S2). Beta-catenin has been suggested to have opposite functions during OL lineage progression (Dai, et al., 2014; Fancy, et al., 2009). Thus, these transcription factors might exert different functions in *Pdgfra+* cells at these developmental stages, most likely depending on which transcriptional complexes they are associated with. These networks of transcription factors involved in patterning and specification/cell commitment were predominant in E13.5, but not at P7 *Pdgfra+/*GFP cells (Figure 2c), when they appear to be replaced by other transcriptional networks leading to the expression of transcription factors such as Sox10, required for OL differentiation (Stolt, et al., 2002). These networks were predicted to be activated by transcriptional regulators such as *Myrf*, *Smad7* and in particular *Tcf7l2* (Figure 2c), which are up-regulated at P7 and have been shown to play important roles in OL differentiation (Bujalka, et al., 2013; Weng, et al., 2012; Fu, et al., 2009; Ye, et al., 2009).

### Single cell RNA-Seq reveals similar transcriptional profiles of post-natal OPCs in the spinal cord and brain

The diversity of biological processes associated with E13.5 *Pdgfra*+/GFP cells (Figure 1f) could reflect a broad cell potential or cell heterogeneity. Since the later would be obscured by bulk RNA-Seq analysis, we performed single-cell RNA-Seq using STRT-Seq technology (Islam, et al., 2014) on 2496 *Pdgfra*+/GFP cells (1514 cells after quality control (QC)) from *Pdgfra*-H2B-GFP (Klinghoffer, et al., 2002) and *Pdgfra*-CreERT-RCE (LoxP-GFP) mice (Roesch, et al., 2008), at E13.5 and P7 (Figure 3a and Figure S1f). For comparison purposes, we also included in the analysis 271 OPCs, 114 committed oligodendrocyte precursors (COPs), and 75 VLMCs from the juvenile and adult CNS (Marques, et al., 2016). In accordance with the bulk RNA-Seq data, there was a clear temporal segregation of E13.5 and P7 cells (Figure 3b). Spatial segregation was also observed, but even to a lesser extent than in the bulk analysis. In fact, subsets of P7 brain and spinal cord cells clustered together in the t-SNE plot, suggesting close similarity (Figure 3b). We performed cell clustering using the BackSpin2 (Marques, et al., 2016) and PAGODA (Fan, et al., 2016) algorithms (Figure S4a,b), both of which revealed similar clusters. In order to establish an unbiased end-point for cluster splitting and avoid oversplitting in PAGODA, we implemented a custom algorithmic threshold, which performs differential gene expression between clusters, and indicates whether they should be merged if no statistical difference in gene expression is detected. BackSPIN2 and PAGODA clusters were merged to form our final cluster set (see Methods). We could identify cells expressing hallmarks of COPs and Newly Formed OLs (NFOL) from the P7 CNS (Figures 3b,c), confirming that these populations appear to arise in the CNS already at P7 and remain transcriptionally similar in the juvenile and adult CNS (Marques, et al., 2016). COPs and NFOLs were found in higher proportions in the spinal cord at P7, highlighting that the process of differentiation is further advanced in this region (Figures 3b, c).

**Fig. 3.**
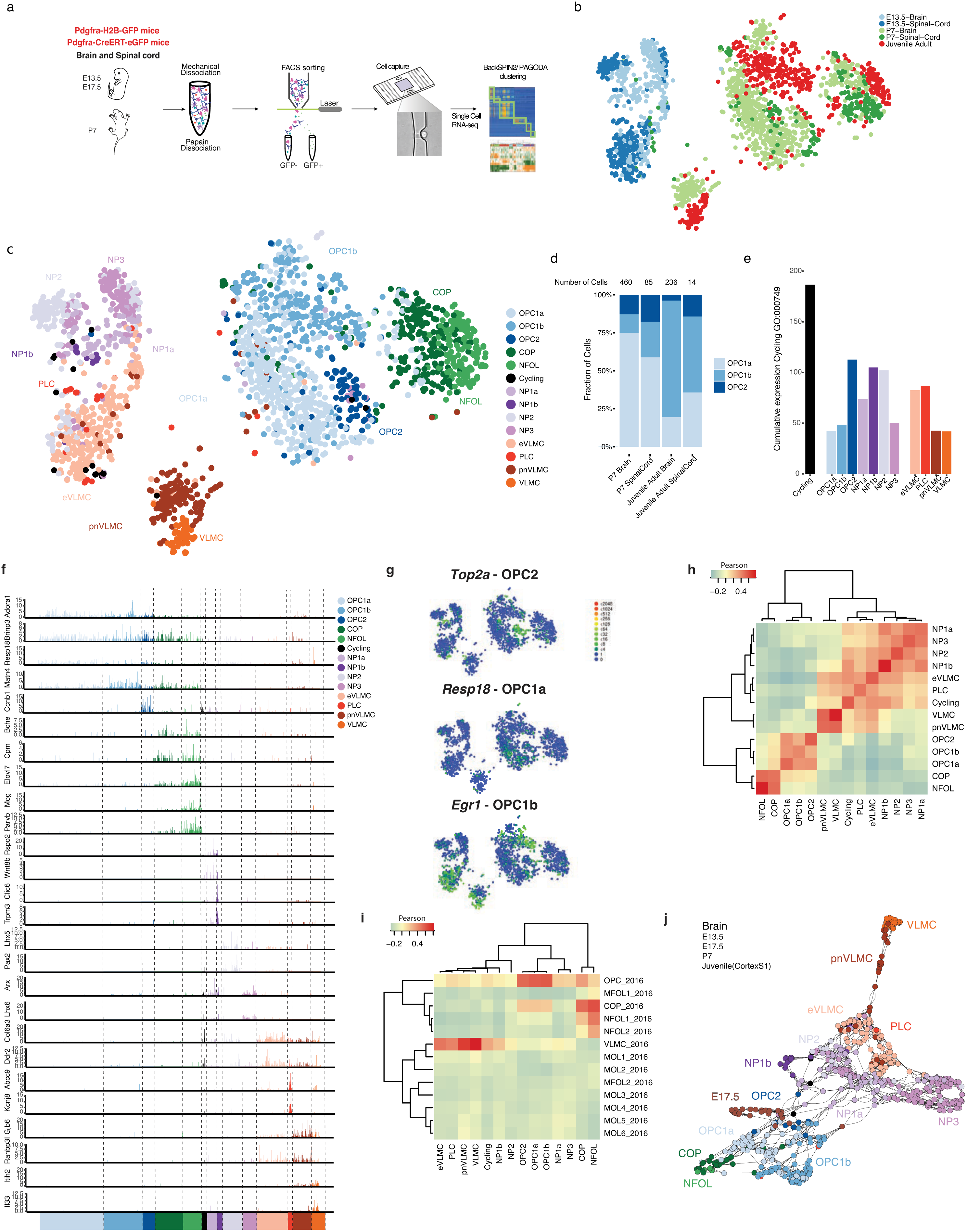
Single cell RNA-Seq reveals similar transcriptional profiles of OPCs in the postnatal spinal cord and brain.

a) Scheme of OPC purification for single cell RNA-Seq;
b-c) t-SNE of single-cell RNA-Seq of E13.5, P7 Pdgfra+/GFP cells, and P20-30, and P60 cells from Marques et. al., highlighting age and region (b) and
c) identified clusters.
d) Fraction of OPC1a, 1b and 2 in each P7, juvenile and adult tissues;
e) Total molecule count of cell cycle gene expression taken from GO:000749 among the different cell populations; f-g) Single cell expression of the most relevant marker genes in all identified populations;
h-i) Hierarchical clustering and correlation analysis of the first 20 principal components of all cells against each other averaged by population (h) and of all the populations when compared with OL
lineage subpopulations identified in Marques et al 2016 (i);
j) SCN3E network analysis of forebrain E13.5, E17.5 and P7 Pdgfra+/GFP cells, with juvenile/adult OPCs, COPs and VLMCs.

Three clusters of P7 and juvenile/adult cells presented markers of OPCs (OPC1a, 1b and 2), (Figures 3c,d). Surprisingly, P7 OPCs from spinal cord and brain clustered together (Figures 3b,c). We performed a removal of hidden confounding factors using the f-scLVM package (Buettner, et al., 2016), which revealed a even more homogeneous distribution of spinal cord and brain derived OPCs (Figure S4c). Batch corrected SCDE differential expression analysis of the single cell populations in the P7 brain and spinal cord derived OPCs failed to identify significant differences in gene expression (not shown). These results highlight the similarity between P7 OPCs from different anterior-posterior regions and indicate that the difference in myelination genes in the bulk population (Figure 1b) is due to the presence of COPs and NFOLs in the *Pdgfra*/GFP+ fraction (Figures 3b, c) and not to intrinsic differences in myelinating potential between OPCs. Furthermore, in combination with our IPA analysis (Figure 2), they suggest a convergence in OPCs cell states in the different anterior-posterior regions.

The three OPC clusters presented unique temporal distributions, with none being present in the E13.5 (Figure 3b,c), OPC1a enriched at P7, and OPC1b at juvenile/adult CNS (Figure 3d). OPC2 was segregated from OPC1a/b and expressed genes involved in mitosis and cell division (Figure 3e, Figure S5a, Table S3). OPC1a and OPC1b clusters were quite similar in their transcriptional profile, which could indicate that they might constitute cell states rather then cell types, or even constitute a single cluster (Figure S5a). However, OPC1a presented distinctive expression of *Resp18* (Figure 3f,g) and genes involved mainly in diverse metabolic processes (mitochondrial energy production, RNA processing), such as ATP synthase, Cox and ribosomal genes (Table S3). OPC1b was characterized by the expression of genes involved in nervous system development and transcription regulation. We also found enriched expression of immediate early genes such as *Fosb*, *Fos*, *Jun*, *and Egr1* (Figure 3g), which might be associated with specific activation of OPC1b cells by the neighbouring neuronal network, although it could also simply reflect cellular stress during the cell extraction procedure. All three OPC cell states were observed in both brain and spinal cord (Figure 3d), highlighting the similarity of OPCs in the anterior-posterior axis of the CNS.

### E13.5 *Pdgfra*+/GFP cells do not have the hallmarks of OPCs

Clustering and differential gene expression analysis allowed the identification of six distinct populations of E13.5 *Pdgfra*+/GFP cells (Figure 3c, Table S3). Four populations (neural progenitors (NP) 1a, 1b, 2 and 3) presented high correlation between themselves (Figures 3h, i). We examined the expression profiles of identified markers of these populations in the E13.5 CNS through publically available *in situ* hybridization data from the Allen Institute for Brain Science (©2008 Allen Institute for Brain Science. Allen Developing Mouse Brain Atlas Available from http://developingmouse.brain-map.org). The NP markers did not present overlapping expression patterns (Figure S5b), indicating that they are indeed expressed in distinct cell populations. NP1-3 expressed markers of the neural progenitor, neuroblast or radial glia lineages (Figure S5a). Indeed, correlation analysis with the embryonic midbrain cell populations identified by single-cell RNA-Seq (La Manno, et al., 2016) indicated a partial similarity between NPs and neuroblast and neuronal progenitor populations (Figure S6a). While some cells within the NP populations expressed individual markers of OPCs (Figure 3f), RNA levels for these genes were relatively low in the cells where they were observed (Figure S5a). Two additional E13.5 clusters expressed higher levels of *Pdgfra* and *Csgp4* (NG2), albeit lower than P7 OPCs (Figure S5a). They also expressed genes involved in collagen formation, such as *Collal*. Correlation analysis with previous published single cell datasets (Marques, et al., 2016; Zeisel, et al., 2015) indicates that these cells are related to pericytes and VLMCs (Figure 3i) and as such they were named embryonic VLMCs (eVLMCs) and pericyte lineage cells (PLCs). Thus, the pericyte/VLMC signature of E13.5 *Pdgfra*+/GFP cells found in the bulk RNA-seq is most likely due to cell heterogeneity than broader cell potential.

### NP1a constitutes a pre-OPC neural progenitor population

NP1a and NP3 presented higher correlation to OPCs than other NPs (Figures 3h,i), and as such, might be the embryonic progenitors of the OL lineage. In particular, a subset of cells of NP1a expressed genes as *Olig1/2* and *Ptptrzl*, but also Nestin (*Nes*), suggesting that they might constitute pre-OPCs. In order to determine how these populations were related, we developed a new algorithm for single-cell near-neighbour network embedding (SCN3E, see Methods). By examining the neighbours for each individual cell, we can infer which cells are most likely to be the progeny or progenitor of others. As a proof of concept, we analysed the OL lineage populations identified in our previous single cell RNA-Seq work (Marques, et al., 2016). SCN3E led to a similar ordering to the one obtained by t-SNE and Monocle, with a clear path from OPCs to MFOLs (Marques, et al., 2016), but allowed a better resolution in the late differentiation/maturation, indicating a branching event at the MFOL stage (Figure S6b). SCN3E analysis for forebrain (Figure 3j) and spinal cord (Figure S6c) for the current dataset indicated that eVLMCs are related to PLCs, while having a weak connection to pnVLMCs and VLMCs in the brain. In accordance to the correlation analysis, NP1a was more related to post-natal OPCs in the brain, which then give rise to COPs and the remaining OL lineage. NP1a (primarily found in the brain and scarce in the spinal cord) and NP2 cells (which are scarce in the brain) are closer to OPCs in the spinal cord SCN3E analysis. As such, we are not able to clear identify which of these populations is progenitor of OPCs in the spinal cord and additional single cell transcriptomic analysis of spinal cord cells would be required. In sum, SCN3E analysis suggests that NP1a might be the progenitors of OPCs in the brain, constituting a pre-OPC cell state in this CNS region

Since there is a clear transcriptional leap between E13.5 progenitors and P7 OPCs (Figure 3c), we performed additional single cell RNA-Seq with E17.5 *Pdgfra*+/GFP cells of Pdgfra-H2BGFP knock-in mice. At this stage, *Pdgfra*+/GFP cells were mainly associated with OPC clusters, although a subset of cells clustered close to E13.5 NP1a cells (Figure S6d). Neural progenitors are nearly absent within the *Pdgfra*+/GFP population, having already given rise to OPCs. We also performed SCN3E analysis of brain single-cell RNA-Seq data, including E17.5 data (Figure 3j). E17.5 OPCs were clustered between E13.5 NP1a cells and P7 OPCs, supporting NP1a cells as pre-OPCs.

### E13.5 *Pdgfra+* cells give rise mainly to OPCs in the postnatal CNS, but also to cells of the pericyte lineage

E13.5 *Pdgfra*+/GFP cells were enriched in genes involved in extracellular matrix and collagen organization, basal lamina and angiogenesis (Figure 1f, Figure S2c) and single cell data indicated the presence of cells of the pericyte lineage. To investigate whether embryonic *Pdgfra+* cells can give rise to cells of the pericyte lineage, we performed lineage-tracing experiments by injecting tamoxifen in E12.5 Pdgfra-Cre^ERT^-LoxP-eGFP mice (Kang, et al., 2010) and examined the GFP+ population at P21 (Figure 4a). While the progeny of the first OPC wave has been reported to be virtually eliminated after P10 (Tripathi, et al., 2011; Kessaris, et al., 2006), we found that at P21 there was still a considerable contribution of E13.5 *Pdgfra+* cells to the OL lineage, and in particular to OPCs (Figures 4b,c and e). 47±6% of the GFP+ cells in the corpus callosum were OPCs (Pdgfra+/Col1-) (Figure 4e), while 15±4% of E13.5 expressed Col1a1, a marker of the vascular and leptomeningeal cells (VLMC) and pericyte lineages (Marques, et al., 2016) (Figure 4d,e). Some of the remaining GFP+ populations were mature OLs (Figure 4c). Similar proportions were observed in the dorsal horn (spinal cord) (Figures 4b,c,e). Thus, a subset of E13.5 *Pdgfra*+/GFP cells (most likely eVLMCs or PLCs) belong to the pericyte lineage and retain this properties or give rise to new cells of this lineage (e.g., pnVLMCs, VLMCs) at postnatal stages.

**Fig. 4.**
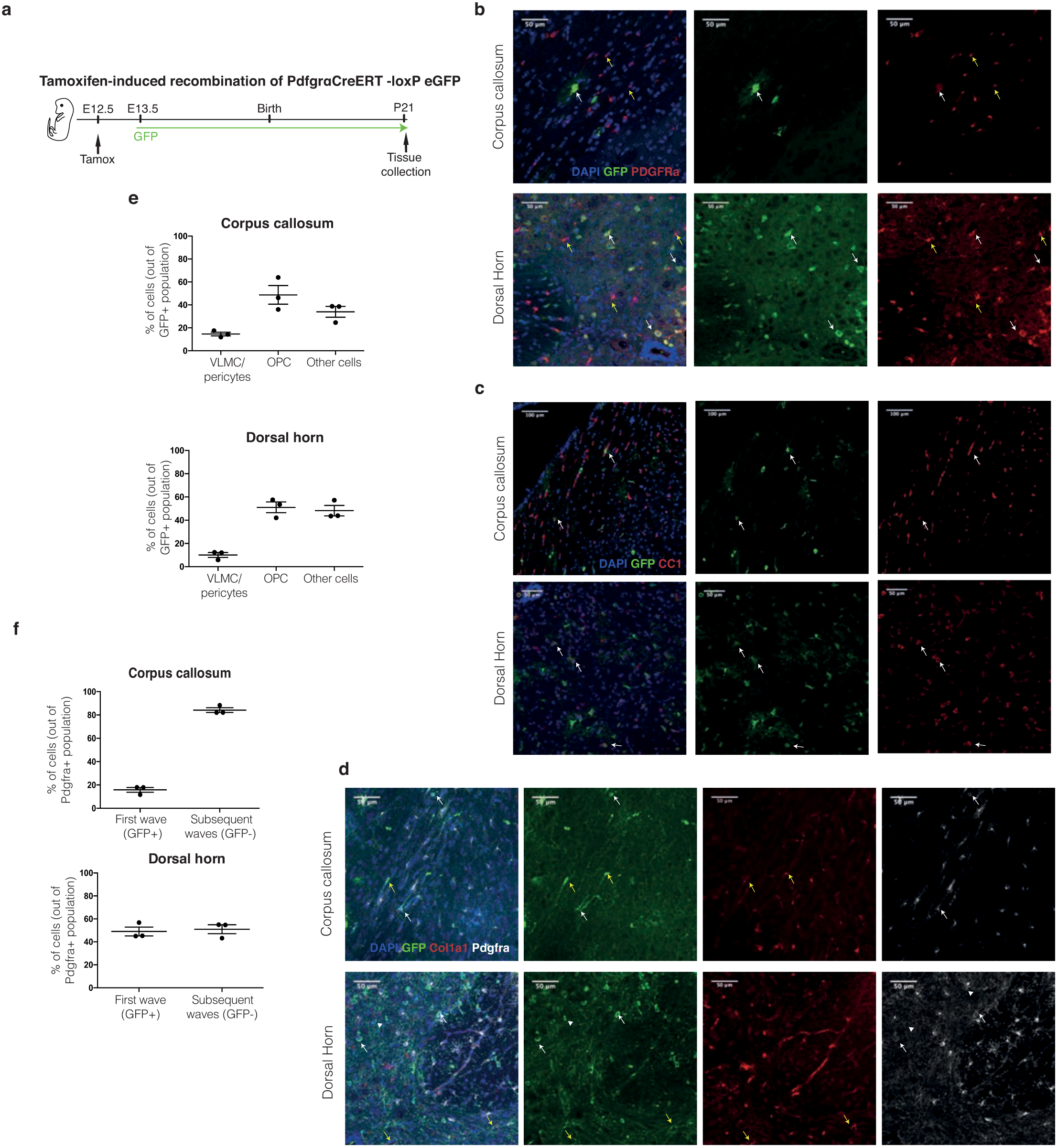
E13.5 Pdgfra+ cells give rise to OPCs and cells of the pericyte lineage.

a) Scheme of lineage tracing experiments in Pdgfra-CreERT-RCE mice; E13.5 Pdgfra+ cells progeny was followed until P21 by GFP expression;
b-d) Immunohistochemistry targeting PDGFRA+ cells (b,d), CC1+ mature oligodendrocytes (c) and cells of the pericyte lineage (Col1a1+, d) in E13.5 Pdgfra+ progeny (GFP+) in corpus callosum and spinal cord (dorsal horn);
e) Quantification of OPCs, VLMCs/pericytes and other cells derived from E13.5 Pdgfra+ cells in the corpus callosum and dorsal horn;
f) Quantification of OPCs derived from the first wave (GFP+/ PDGFRa+) and second/third waves (GFP-/ PDGFRa+) in the corpus callosum and dorsal horn.

Since our data suggested that the first wave of oligodendrogenesis gives rise primarily to OPCs after birth, we examined the percentage of OPCs (Pdgfra+/Col1-) originating from this wave (GFP+ cells), compared to subsequent waves (GFP- cells). In the corpus callosum, less than 20% of OPCs were derived from the first wave, while the majority was derived from subsequent waves (Figure 4f). This is consistent with the findings that 20% of OL lineage cells (Sox10+) in the corpus callosum at P12-P13 are derived from the first/second wave, while 80% originate from the third one (Tripathi, et al., 2011). In contrast, in the dorsal horn 48±4% of the OPCs were derived from the E13.5 wave (Figure 4f). Thus, our results indicate that the progeny of first wave of E13.5 *Pdgfra+* progenitors persists in the juvenile CNS, giving rise not only to a specific population of OPCs, but also to cells of other lineages, such as pericytes.

### Rewiring of transcriptional networks during the transition between pre-OPCs and OPCs

In order to define functional heterogeneity within the single-cell dataset and to reveal putative regulators acting at the different time points within our dataset, we performed weighted correlation network analysis (WGCNA, see Methods). We identified 10 robust gene modules, each correlating with expression profiles of different cells with few overlaps (Figure 5a,b and Tables S4, S5). The largest gene module corresponded to OPCs and to genes involved in regulation of synapse structure or activity, and ion channel activity. Genes associated with this module included the classical oligodendrogenesis transcription factors *Olig1*, *Olig2*, *Sox10*, as well as genes such as *Ptprz1*, *Cspg4*, and *Pcdh15*. The OPC2 population correlated well with a cell cycle gene module, involving regulators such as *Hmgb2* and *Cenpa*, and highly associated genes such as *Top2a*, *Birc5*, *Mki67*. COPs and NFOLs correlated with a module enriched for myelination genes, with highly intramodularly connected regulators such as *Tcf7l2* and *Zeb2*, and genes such as *Cnp*, *Mbp*, *Plp1*, *Itpr2* and *Qk*. VLMCs, pnVLMCs, eVLMCs and PLCs correlated with gene modules involved in vascular development, extracellular matrix formation and smooth muscle contraction. Three gene modules were correlated with the NP populations. NP3 was mainly associated with a neuron commitment module characterized by expression of regulators such as *Arx*, *Dlx6os1*, *Dlx1/2/5 and Lhx6*. NP2, NP1a and a small subset of eVLMC cells correlated with a gene module with predicted regulators such as *Sox11*, *Rbfox2*, *Hoxb3*. Other genes associated with these modules were *Cd24a*, *Tubb3*, and *Stmn1/2*. NP1b was correlated with a small gene module specific for epithelium development and cilium organization.

**Fig. 5:**
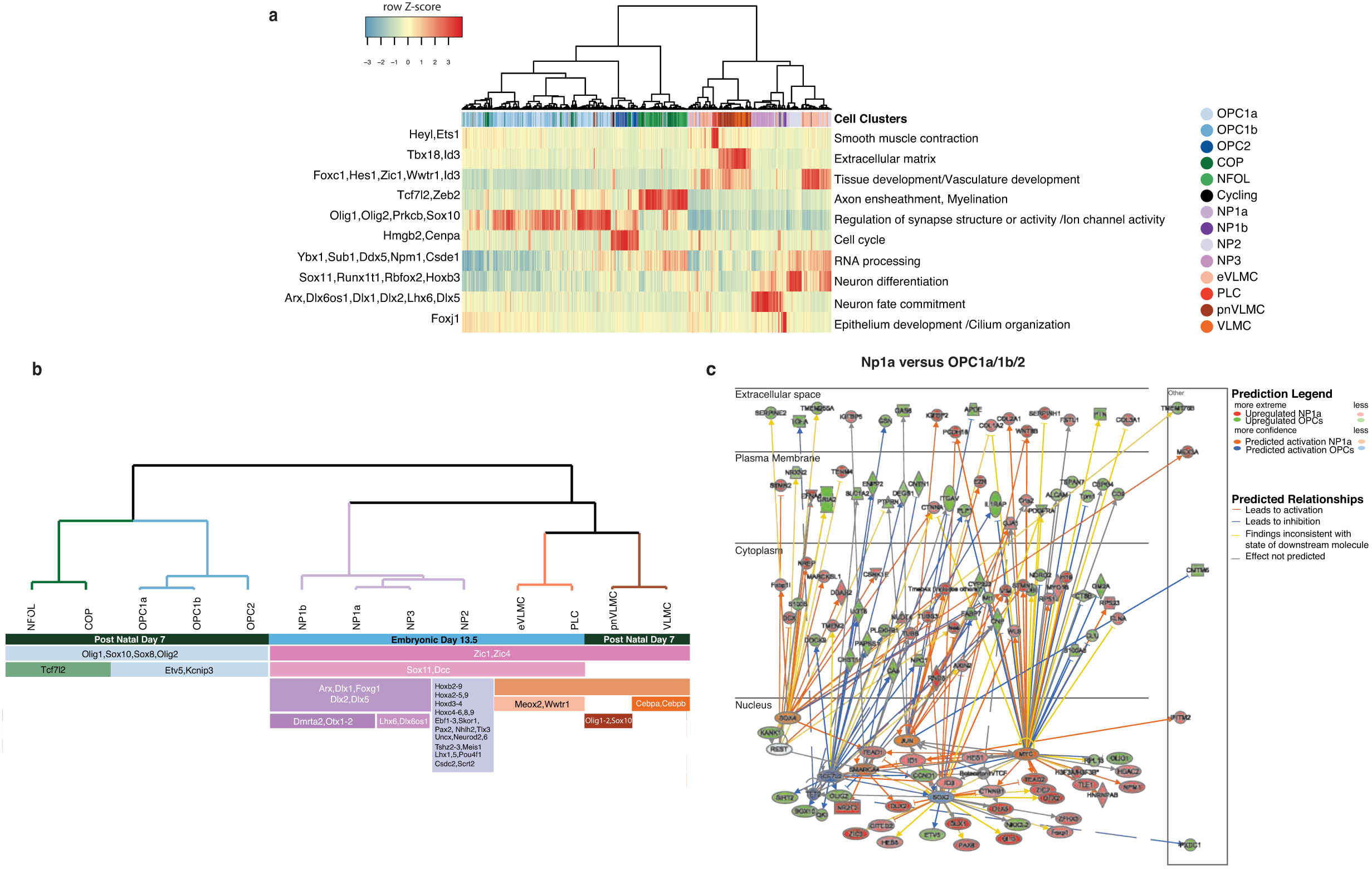
Transcriptional rewiring of OL progenitors into postnatal OPCs.

a. WGCNA heatmap showing the row Z-score for each gene module;
b. Overview of transcription factors associated with specific populations as defined by SCDE dendogram comparison analysis;
c. Network of genes controlled by the IPA- identified upstream regulators, including Cttnb1, Jun, Myc, Rest, Smarca4, Sox2, Tcf7l2, Tet2, and Sox4 in NP1a and all OPC single-cell RNA-seq datasets;

We further analyzed the transcriptional shifts between pre-OPCs (NP1a) and OPCs using IPA (Figure 5c). In pre-OPCs, transcription factors involved in patterning, negative regulators of differentiation or cell specification and commitment were predicted to be regulated in particular by *Myc and Sox4*, but also by beta-catenin (*Ctnnb1*), *Jun* and the chromatin remodeller *Smarca4*. In contrast, *Sox2*, while enriched in pre-OPCs in comparison to OPCs (Figure 5c), was predicted to be active in OPCs. Sox2 was reported to prevent OPC specification (Kondo and Raff, 2004), but also to be necessary for OPC proliferation and differentiation (Zhao, et al., 2015; Hoffmann, et al., 2014), highlighting that the activity of many of the identified transcriptional regulators is indeed context-dependent. OPC transcriptional networks were predicted to be activated by *Tcf7l2*, which is in accordance with the IPA analysis of the bulk RNA-Seq and by WGNA analysis (Figure 5a,b). *Bmi1* (*Commd3-Bmi1*) and *Kat6*, identified in the IPA analysis of the bulk RNA-Seq data (Figure 2), were also predicted to be active in NP3 and NP2, respectively (Table S6). We also performed IPA to investigate the segregation between the three identified OPC states (Figure S6 and Table S6). OPC2 is likely to represent a proliferative OPC state. The forkhead transcription factor *Foxm1* and *Myc* were predicted to be activated in the transition between OPC1b-OPC2 and OPC1a-OPC2, being upstream several cell cycle related genes, such as *Ccna2* (cyclin A2), *Cdk1*, *Cdc20*, among others. *Foxm1* is indeed expressed in higher levels in OPC2, unlike *Myc*. In addition, *Tp53, Pten and* the histone lysine-specific demethylase 5b (*Kdm5b*) were predicted to be active in OPC1a/b and inhibit genes that are enriched in OPC2, such as *Birc5*, *Aurka*, *Aurkb*, involved in mitotic cell cycle processes.

### Novel transcripts and non-coding RNAs regulating transitions between progenitor states within the oligodendrocyte lineage

Non-coding RNA (ncRNAs) have recently emerged as master regulators of epigenetic landscapes and developmental processes (Batista and Chang, 2013), and could thus be potential regulators of the identified modules and cell populations. As we used polyA-neutral stranded bulk RNA-Seq, we were able to analyse and identify the differential expression of annotated and novel non-coding and protein-coding RNAs in *Pdgfra+/GFP* cells (Figure S7a,b). A total of 3,653 putative non-coding and 1996 putative coding novel multiexonic transcripts were predicted, of which 1853 were differentially expressed (Table S1). 938 annotated ncRNA were also differentially regulated in the bulk data (Figure 6a). Furthermore, a subset of the novel and annotated non-coding transcripts, were found to be specific for the populations identified in the single cell RNA-Seq (Figure 6b). In the bulk data, precursor transcripts of miR-219 and miR-124 were specific for the postnatal spinal cord and embryonic samples, respectively (Figure 6a), in line with their reported roles in the transition from proliferating OPCs to myelinating OLs (miR-219) (Dugas, et al., 2010) and neural differentiation (miR-124) (Maiorano and Mallamaci, 2010). The *Rmst* lincRNA, required for neurogenesis (Ng, et al., 2013), was among the most strongly down-regulated lincRNA in both brain and spinal cord during development. Accordingly, it was a marker for the pre-OPC (NP1a) and NP2 neuronal populations. Other transcripts specific for the NP populations and downregulated in the bulk data during development included *Dlx1as* (NP1a, NP3), *Dlx6os1* (NPs, NP1a, NP1b, NP3); *Lhx1os* (NP2), and the novel assembled transcripts *Ept1-au1* (MSTRG.27163) and *Scaper-au1* (MSTRG.37353), both specific for the NP1b population. The imprinted lncRNAs *Airn* and *H19* were also both strongly downregulated during development, and revealed to be specific for the eVLMC population.

**Fig. 6.**
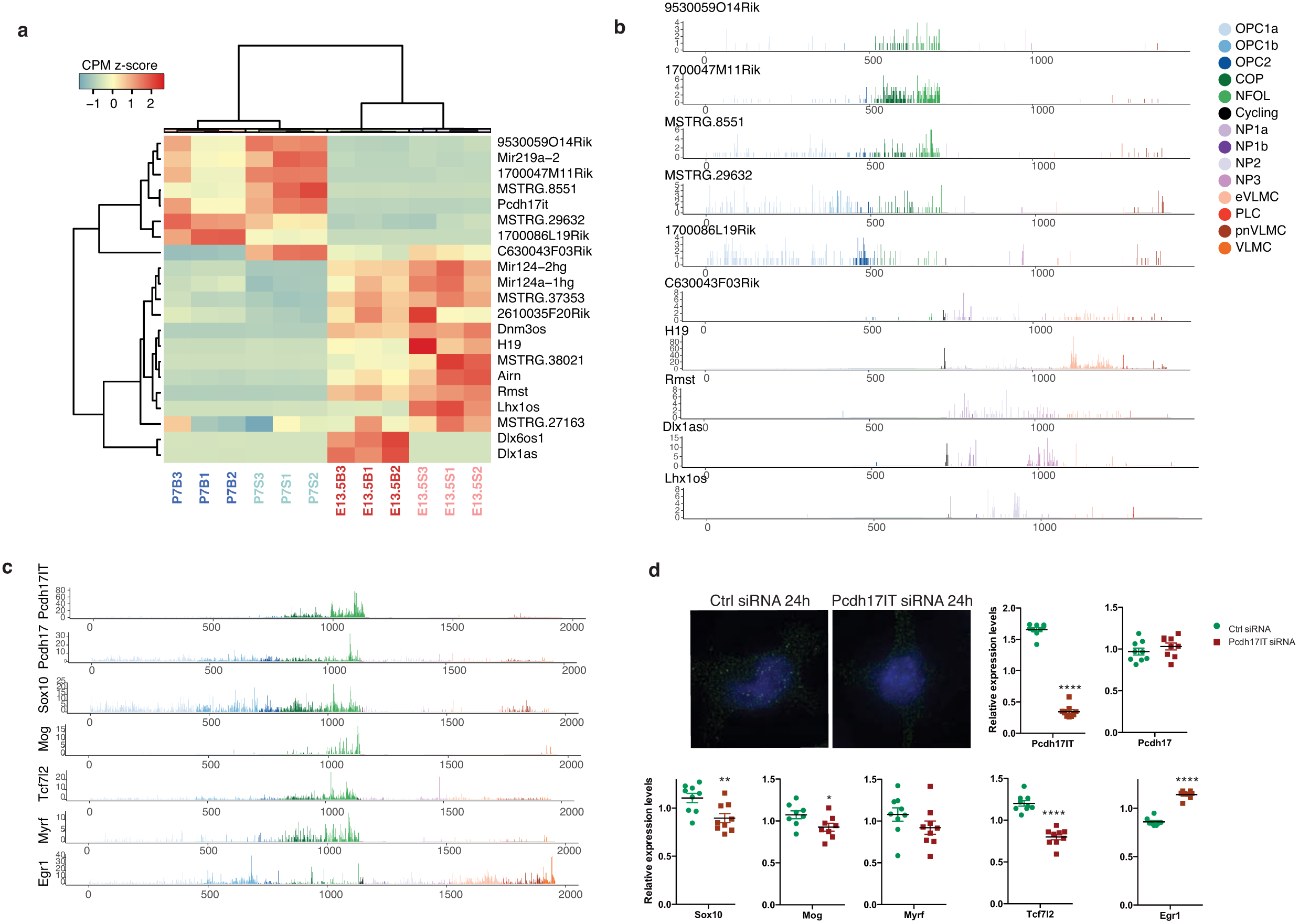
Novel transcripts and non-coding RNAs regulating transitions between progenitor states within the oligodendrocyte lineage.

a. Novel transcripts and annotated lncRNAs differentially expressed in bulk data;
b. Expression in the different single cell clusters of the novel transcripts and annotated lncRNAs identified in bulk data;
c. Expression levels of Pcdh17IT, its neighbouring gene, OL markers and TF co-expressed in the same populations (COP/NFOL) or in the transition to these populations;
d. Analysis of knockdown of lncRNA Pcdh17IT in Oli-neu cell line by smFISH (high magnification of images shown in Fig. S7) and qRT-PCR (n=8-9); two-way student t-test; p- value: * < 0.05, ** <0.01, ***<0.001, ****<0.0001;

Many of the transcripts that were upregulated during development in the bulk samples were specific for the early stages of OL differentiation. This included *1700047M11Rik* (COP, NFOL, and OL), *9530059O14Rik* (COP and NFOL), and two novel assembled transcripts - *MSTRG.8551* (COP, NFOL and OLs) and *MSTRG.29632* (OPC1b) (Figure 6b). We also identified the long non-coding RNA *9630013A20Rik* (also referred as *LncOL1* (He, 2016)) to follow this pattern and to be specific for the COP and NFOL populations (Figure 6c). Since it is localized in the intron of the gene *Pcdh17*, and following HUGO Gene Nomenclature Committee guidelines (Wright, 2014), we named it *Pcdh17IT*. In line with recent results (He, 2016), we found that knocking down this ncRNA in a mouse oligodendrocyte precursor cell line (Figure 6d) led to downregulation of *Tcf7l2* and *Myrf* (although in lower extent), both transcription factors specifically expressed in COP and NFOL and that are key hubs in the postnatal transcriptional networks (Figures 2 and 5), as well as *Sox10* and *Mog* (Figures 6c, d). In contrast, *Pcdh17IT* knockdown led to an increase of *Egr1*, which is mainly expressed in OPC1b and downregulated upon the transition to COPs (Figure 6c, d). Thus, these results indicate that ncRNAs are specifically expressed in distinct OL progenitor populations and their progeny, and suggest that they regulate transitions between these states by regulating key transcription factors.

### E13.5-derived post-natal OPCs and OPCs derived from subsequent waves have similar transcriptional profiles and electrophysiological properties

Our single cell transcriptomics analysis indicates that embryonic OL progenitors undergo a process of rewiring of their transcriptional networks, converging in postnatal OPCs states equipped for receiving inputs from the neighbouring neuronal network and enabling them to undergo differentiation. In order to determine whether this convergence occurs during development *in vivo*, we performed lineage tracing of *Pdgfra+*cells originating from only the first wave (tamoxifen treatment at E12.5) or from all waves (tamoxifen treatment at P3), and assessed their progeny at P7 by single cell RNA-Seq (Figure 7a). We observed that OPCs derived from E13.5 or from subsequent waves intermingled (Figure 7b), with OPC1a/b and OPC2 cells being progeny of both the first wave and the subsequent waves (Figure 7c). OPC1b had a smaller representation in P3-derived OPCs, which might suggest that a defined temporal window is required for the maturation of *Pdgfra+*/GFP into OPC1b, as well as pnVLMCs and COPs. We also analysed the lineage-traced cells focusing only on the expression profiles of genes involved in electrophysiological activity, specifically glutamate receptor, potassium channel, voltage gated ion channel, and GABA receptor genes. t-SNE representation indicated that variation between these genes could not segregate txE12.5 from txP3 cells (Figure 7d). Accordingly, these genes were expressed in a stochastic manner in both populations (Figure 7e). Thus, our results indicate that OPCs from different development temporal and spatial origins indeed converge into three transcriptional states at post-natal stages.

**Fig. 7.**
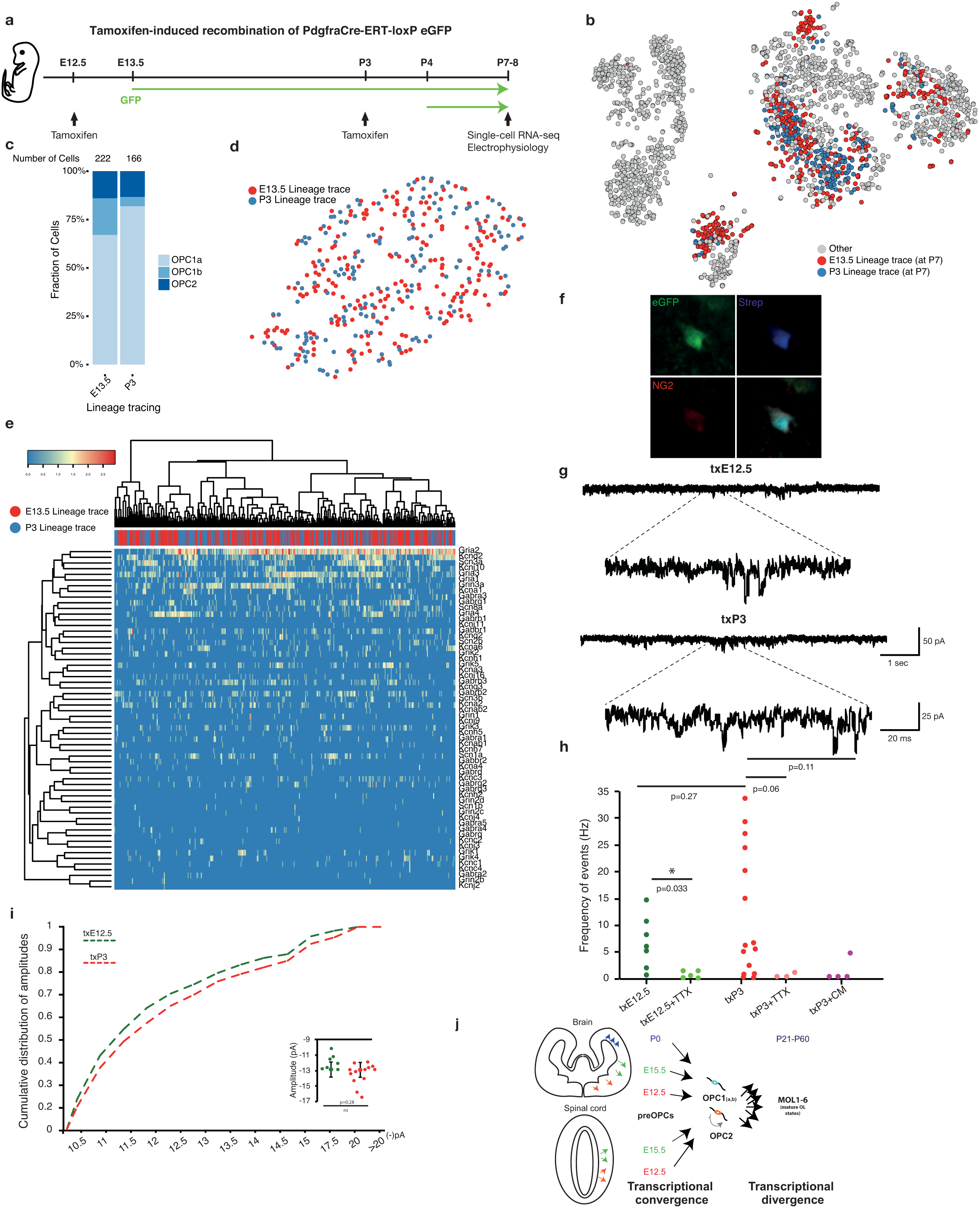
Similar single-cell transcriptomic and electrophysiological profiles of cells derived from the first and subsequent waves of oligodendrogenesis.

Scheme of lineage tracing experiments in Pdgfra-CreERT-RCE mice; E12.5-13.5 and P3-5 Pdgfra+ cells progeny was identified by GFP expression at P7-8 when single cell RNA-Seq and electrophysiological recordings were performed; b) t-SNE illustrating GFP+ cells from the lineage tracing of E12-5-13.5 or P3-P5 Pdgfra+ cells in the PdgfraCreERT mice at P7; c) Fraction of OPC1a, 1b and 2 in each lineage tracing experiment; d) tSNE clustered using glutamate receptor, potassium channel, voltage gated ion channel, and GABA receptor genes illustrating homogeneous distribution of E13.5 and P3 lineage traced OPCs; e) Hierarchically clustered heatmap showing the expression of glutamate receptor, potassium channel, voltage-gated ion channel, and GABA receptor genes in E13.5 and P3 lineage traced OPCs; f) Representative image showing a recorded cell labelled with biocytin-streptavidin (blue) and expression of NG2 (red); g) Representative voltage-clamp traces of OPCs (held at -70 mV) showing inward spontaneous postsynaptic currents; d) Frequency of events in seven txE12.5-cells (dark green) and fifteen txP3-cells (red). Tetrodotoxin (TTX) strongly reduced frequency spontaneous of events, shown as txE12.5+TTX (five cells) and txP3+TTX (three cells). Blockage of glutamatergic receptors with CNQX/MK801 showed similar effect (txP3-CM, four cells); h) Cumulative distribution of events’ amplitudes and on the right side insert with average amplitudes for each cell. P-values correspond to two-tailed Student’s t-test for independent (between groups) or paired-samples (for pharmacology experiments). j) Model for transcriptional convergence of the different waves of progenitors of oligodendrocyte lineage cells.

Our single-cell RNA-Seq experiments coupled with lineage tracing would suggest that E13.5 derived OPCs at P7 have similar electrophysiological properties to OPCs derived from subsequent waves. In order to investigate if this is indeed the case, we performed whole-cell voltage-clamp recordings of corpus callosum OPCs at P7-P8 derived from either the first wave (tamoxifen treatment at E12.5 – txE12.5) or all waves (tamoxifen treatment at P3 – txP3) (Figures 7a,f). The two groups (slices from 2 animals txE12.5 and from 4 animals txP3) exhibited comparable passive intrinsic properties: input resistance (txE12.5 1.6 ± 0.2 GΩ; txP3 1.6 ± 0.2 GΩ; (n) of cells = 6 and 15 respectively) and membrane capacitance (txE12.5: 34.3 ± 8.7 pF, n=7; txP3: 30.7 ± 6.1 pF, n=15). Recorded events had similar average amplitudes (txE12.5: -12.28 ± 0.39 pA, n=7; txP3: -13.0 ± 0.4 pA, n=15). The frequency of events in the two populations was also not significantly different (Figures 7g-i). However, while txE12.5 cells received less frequent events in a more uniform manner (6 ± 1.89 Hz, n=7), txP3 cells exhibited a larger heterogeneity, with 30% of the cells having more frequent events (11.4 ± 3.1 Hz, n=15; p=0.3). Concurrently, the cumulative distribution of event amplitudes displayed somewhat larger events recorded on OPCs derived from txP3 (Figure 7i). In order to confirm that these events are due to spiking activity in neighboring neurons, we blocked excitation by applying 10μM of Tetrodotoxin (TTX) in slices from 2 animals per time point. This drastically reduced the frequency of events in both groups (Figure 8c): txE12+TTX (n=5; 0.93 ± 0.33 Hz; p=0.038) and txP3+TTX (n=3; 0.54 ± 0.32 Hz; p=0.075). Accordingly, when glutamatergic receptors were blocked, we also observed a robust decrease in number of events (txP3+CM- 1.2 ± 1.3Hz; n=4; p=0.11, Figure 7h). As such, OPCs derived from the E13.5 wave present similar electrophysiological properties to OPCs derived from subsequent waves.

## Discussion

We have previously shown that OL lineage in the juvenile and adult CNS is more heterogeneous than previously anticipated, with several intermediate states after differentiation of OPCs and six matures OL states (Marques, et al., 2016). Here we provide evidence that progenitor cells of the OL lineage in different anterior and posterior regions of the CNS are distinct at E13.5, but converge into three OPC transcriptional states at P7 (Figure 7j). Our results from bulk and single-cell RNA-Seq suggest that embryonic patterning transcription factors, apart from defining region identity, are upstream of a network of inhibitory regulators that are important for the maintenance of the progenitor state. This transcriptional network is replaced at postnatal stages in the anterior and posterior CNS by a convergent transcriptional network, associated with electrophysiological responses and compatible with activity driven differentiation and myelination (Koudelka, et al., 2016; Gautier, et al., 2015; Wake, et al., 2015; Gibson, et al., 2014; Lundgaard, et al., 2013; Demerens, et al., 1996). We identify several cohorts of transcriptional regulators, including transcription factors, chromatin modifiers and novel non-coding RNAs, which are involved in this transcriptional rewiring. Given this convergence into similar OPC states at post-natal stages, when differentiation start, subsequent cell state diversification into six mature OL cell states (Marques, et al., 2016) is thus likely not cell intrinsic, but rather induced by the local environment OPCs are exposed to upon differentiation (Figure 7j).

Previous transcriptomics studies on the oligodendrocyte lineage were based on bulk population expressing *Pdgfra* and focused on single regions, as the forebrain (Moyon, et al., 2015; Zhang, et al., 2014; Cahoy, et al., 2008). These studies were performed with bulk RNA-Seq or microarrays, and suggested for instance that neonatal and adult OPCs presented different transcriptional profiles (Moyon, et al., 2015). In our study, bulk and single-cell RNA transcriptomics were for the first time performed on the exact same *Pdgfra*+ cell population at different stages and regions, giving an unbiased and clear understanding of what exactly these cells are at the molecular level – and how diverse the *Pdgfra*+ population is. Bulk RNA-Seq analysis of FACS-sorted *Pdgfra*+/GFP cells suggests heterogeneity between P7 OPCs from the spinal cord and brain, with the former expressing genes involved in myelination at substantially higher levels. However, scRNA-Seq analysis of the populations present at this stage indicates that OPCs are unexpectedly similar at the transcriptional level, highlighting the strength of single-cell RNA-Seq analysis to deconvolute bulk transcriptomics analysis. OPCs in different anterior-posterior regions of the CNS converge into similar transcriptional profiles at P7 that are compatible to their main functions at that stage (integration of neuronal activity and differentiation). Nevertheless, it is possible that small cohorts of differentially expressed genes might be involved in additional functions of OPCs, contributing to different functional states within the OPC cell type within the different regions or stages.

Progenitor heterogeneity is currently defined by its progeny, the cell types the progenitor has potential to give rise to. These cell types are usually investigated with lineage tracing analysis, which depends on the identity of the original gene used for labeling the progenitor cell, and on the small subset of pre-selected immunohistochemical markers used to identify the progeny cell types. The strength of the transcriptomics approach used in this study is that it allows to assess progenitor heterogeneity in an unbiased manner, by identifying cells with distinct transcriptional profiles. By combining E13.5 transcriptomics data with 1) data acquired in later time points, when the progeny of oligodendrocytes have already arisen, and 2) lineage tracing analysis, we were able to link distinct neural progenitor populations with their potential to give rise to specific cells types. Previous lineage tracing suggested that Pdgfra+ OPCs are already specified at E12-E12.5 (Kessaris, et al., 2006). In contrast, our approach, by considering all genes characteristic of the E13.5 Pdgfra+/GFP cells, allowed the identification of four neural progenitor subpopulations, including a pre-OPC population (NP1a), which have distinct gene expression profiles from OPCs (Figure 3). At E17.5, these neural progenitors are nearly absent within the *Pdgfra*+/GFP population, having already given rise to OPCs (Figure S6).

Our data indicates that only a subset of *Pdgfra+* at E13.5 express markers of the future oligodendrocyte lineage, and that other subsets have transcriptional profiles compatible of progenitors of other cell lineages. Lineage-tracing had previously indicated that a small subset of postnatal *Pdgfra*+ cells can give rise to neurons, although it was not clear whether this was a technical artifact due to spurious Cre recombinant (Clarke, et al., 2012). Motor neurons are specified at E9-10.5 at the ventral CNS from an Olig2 domain, while a subsequent wave of oligodendrogenesis starts occurring at E12.5, with the emergence of *Pdgfra+* cells (Rowitch, 2004). Thus, oligodendrogenesis rather than neurogenesis should be captured from this domain at E13.5. From the NP populations identified, NP2 cells express lower levels of *Pdgfra* than OPCs. It is possible that progenitors of NP2 cells had transiently expressed *Pdgfra* before E13.5, leading to GFP expression. NP2 present transcription factors associated with neuron specification/differentiation, such as *NeuroD1, NeuroD2, NeuroD6* (Figure S5). Interestingly, *Pdgfra* was observed in the dorsal CNS in the areas around this stage, and have been hypothesized to give rise to interneurons (Pringle and Richardson, 1993). NP2 could thus represent this population. Indeed, NP2 markers are found to be expressed in dorsal part of E13.5 dorsal cord (Figure S5b). Since we performed lineage tracing with Pdgfra-Cre-ERT/RCE mice upon injection of tamoxifen at E12.5/13.5 and P3/P4 and combined with single cell RNA-Seq at P7/8, we could investigate whether this population can give rise to neurons. Correlation analysis of this dataset with a previously published single cell RNA-Seq dataset (Zeisel, et al., 2015) indicates that the lineage-traced cells show high correlation with oligodendrocytes and astrocytes/ependymal cells, and much lower with microglia, endothelial-mural or neurons (Figure S7). Nevertheless, four cells of E13.5-derived cells have a high correlation with neurons (Figure S7). Additional single-cell RNA-seq will be required to elucidate whether embryonic *Pdgfra*/GFP+ cells indeed can give rise to neurons.

We observed E13.5 *Pdgfra*+ populations that express pericyte and VLMC markers, and can give rise to cells of the pericyte lineage, an important finding considering the elusive nature of pericytes progenitors (Armulik, et al., 2011). Pericytes and oligodendrocyte progenitors have been described to share antigenic properties, including expression of *Pdgfra* and *Cspg4* (NG2). Adult *Pdgfra*+ cells have also been described to occasionally give rise to pericytes (Kang, et al., 2010). Thus, it is possible that *Pdgfra*+ eVMLC or even one of the NP populations are the progenitors of at least a subset of cells in the pericyte lineage. Interestingly, it has been previously reported that E7.5 Sox10+ neural crest cells give rise to pericytes in the adult CNS (Simon, et al., 2012). While we do not detect Sox10 expression in eVLMCs or NPs, it is possible that these cells are derived from an earlier progenitor population expressing *Sox10*. pnVLMCs, in contrast to pericytes and eVLMC, express several genes that are present in cells of the OL lineage, such as *Plp1, Mbp, Trf*. This could suggest a lineage relationship with oligodendrocytes. Nevertheless our network analysis (Figures 3 and S6) indicates that such a link is unlikely. Lineage tracing using eVLMC or NP specific markers might elucidate whether indeed any of these populations are progenitors within the pericyte lineage.

We found that that the progeny of E13.5 *Pdgfra*+ cells is not eliminated at the juvenile CNS, but rather that these cells can give rise not only to OLs, but to pericytes and to OPCs (Figure 4). OPCs derived from the E13.5 wave presented similar electrophysiological properties to OPCs derived from subsequent waves (Figure 7). Nevertheless, there was a bimodal pattern of electrophysiological activity for P3 derived OPCs. A subset of second/third-wave derived OPCs are more responsive to the neighbouring neuronal network than E13.5 derived OPCs (Figure 7h), suggesting a higher heterogeneity of second/third wave-derived OPCs than first wave-derived OPCs. When analysing the overall transcriptional profile of OPCs derived from the E13.5 wave and other waves, no heterogeneity was found (Figure 7) indicating that transcriptional state could thus not account for the observed electrophysiological differences. We observed scattered expression of several ion channels and glutamate receptors (Figure 7), which can be consistent with cell heterogeneity at an electrophysiological level. Techniques as Patch-Seq (Fuzik, et al., 2016) might be able to address if this indeed the case. Alternatively, other post-transcriptional events, such as asymmetric distribution of transcripts in OPCs processes (Thakurela, et al., 2016) or translational events might account for the observed differences. Diversity of cell states arises early during development in neuronal lineages, with progenitors expressing patterning transcription factors that will ultimately determine the identity of their progeny. Interestingly, many of these transcription factors are actively downregulated upon terminal neuronal differentiation, while their ectopic reactivation has been linked with cell death and degeneration (von Schimmelmann, et al., 2016) Nevertheless, the attenuation of patterning/specification transcription we observe in the oligodendrocyte lineage is not a common event during neural development. For instance, it does not occur within the dopaminergic neuronal lineage, where patterning transcription factors such as Lmx1a/b and Nurr1 continue to be expressed at adult stages and have distinct functions relative to their developmental ones (Doucet-Beaupre, et al., 2016; Kadkhodaei, et al., 2013). Our results indicate that downregulation of patterning and specification transcription factors appear to be required for the establishment of an OPC transcriptional state. Thus, abnormal re-expression of transcription factors expressed during embryogenesis might be deleterious to postnatal OPCs or might prevent their capacity to differentiate in the context of disease.

## Methods

### Animals

Mice line used in this study included Pdgfra-cre-ERT/RCE (Kang, et al., 2010) (see lineage tracing below) and the Pdgfra-H2BGFP knock-in mouse (Klinghoffer, et al., 2002) (background C57BL/6NJ), in which H2B-eGFP fusion gene is expressed under the promoter of the OPC marker, *Pdgfra*. Mice homozygous for this knock-in targeted mutation have an embryonic lethal phenotype, with half of the embryos failing to survive past embryonic day 12.5 and the remainder failing to survive beyond embryonic day 15.5 (https://www.jax.org/strain/007669). Therefore, we used heterozygote mice, in which *Pdgfra* is expressed mainly in OPCs but also in some extent in the early stages of OL differentiation, due to GFP half-life (Clarke, et al., 2012). Mice were time mated to obtain embryos with 13.5 days or post-natal day 7 pups. Gender was randomized since the experiments were mainly done in embryos and pups. The following light/dark cycle was used: dawn 6.00-7.00; daylight 07.00-18.00; dusk 18-00-19.00; night 19.00-06.00. A maximum of 5 adult mice per IVC-cage of type II Allentown. Breedings were done with 1 male and up to 2 females. All experimental procedures performed followed the guidelines and recommendations of local animal protection legislation and were approved by the local committee for ethical experiments on laboratory animals (Stockholms Norra Djurförsöksetiska nämnd in Sweden).

### OPC extraction

Embryos with 13.5 days and pups from post-natal day 7, from both genders of the Pdgfra-GFP mice line were used to extract OPCs. The dissociation method varied depending on the stage. For embryonic stage, forebrain and spinal cord were excised and tissue was mechanical dissociated in HBSS with ions using 3 different Pasteur pipettes of decreasing diameter. For post-natal stages, the same tissues were dissociated with the Papain Neural dissociation kit from Miltenyi, following the manufacturer’s instructions.

### FACS sorting

The single cell suspension from embryonic and post-natal tissue was FACS sorted for GFP cells using a BD FACSAria III Cell Sorter B5/R3/V3 system. For bulk sequencing, cells were collected in non-sticky RNAse free tubes containing RNA Later, Qiazol was added and samples were snap frozen until further RNA extraction. For this, pooling of mice samples was performed to obtain at least 50000 cells. For single cell, cells were collected in cutting solution with 1% BSA and quickly prepared for capture on the C1 fluidigm system. FACS gating strategy is presented in Figure S1. We observed a gradient of GFP+ cells at P7. This gradient was also previously observed in other studies with the Pdgfra-H2B-GFP mice, where the OPC population was selected for the expression of high levels of GFP (Moyon, et al., 2015). We performed qRT-PCR studies and determined that while the GFP++ population expressed *Pdgfra*, *Olig2* and *Plp1*, the GFP+ population lacked expression of these markers (data not shown, n=1). Since the main focus of our manuscript is OPCs, which express *Pdgfra*, we did not proceed to the analysis of the GFP+/Pdgfra-population.

### RNA extraction for Total RNA sequencing

RNA was extracted with miRNeasy micro kit from Qiagen, following manufacturer´s instructions with minor modifications. Briefly, qiazol extracts were thawed and vortexed for 1min. Chloroform was applied and mixture was transferred to a MaxTract High density column. From that point on, the extraction was then performed according to the kit´s manual for low input samples. RNA was then measured with QuBit RNA HS assay kit and Agilent RNA 6000 Pico Kit. 50 ng of RNA from each E13.5/ P7 sample was used for library preparation.

Each sample represented a pool of *Pdgfra+* cells collected from 4 – 32 animals to achieve the 50 ng required for library preparation.

### Library preparation for Total RNA sequencing

Illumina’s TruSeq Stranded Total RNA Library Prep Kit was used according to manufacturer’s instructions. Three pooled biological replicates were sequenced for each developmental stage and brain region, and a total of 38 - 66 million 150 bp strand-specific paired-end reads were generated for each replicate, comprising a total of 620.5 million reads across all datasets (Figure S1).

### Statistical analysis of gene expression

After read quality control using FastQC (Andrews, 2010), STAR (Dobin, et al., 2013) (v.2.5.0a) was used to map reads to the mouse mm10 genome. GENCODE (Mudge and Harrow, 2015) M8 annotations were used to construct the splice junction database and as a reference for the count tables. Gene-level count tables were obtained using featureCounts (Liao, et al., 2014) v1.5.0-p1, assigning multi-mapping reads fractionally to their corresponding loci and using convergent rounding to convert the resulting count tables to the nearest integer values. Bioconductor packages were used for data processing and analysis (Gentleman, et al., 2004), and the biomaRt library was used for querying annotations and mapping across gene identifiers (Durinck, et al., 2009). The limma package was used for differential gene expression analysis (Ritchie, et al., 2015), with normalisation carried out using the voom approach (Law, et al., 2014). Genes were tested for differential expression if they displayed 0.7 counts per million in at least three of the libraries, and considered differentially expressed if found to have a Benjamini-Hochberg adjusted p-value < 0.05 and a greater than twofold change in expression between queried samples. Exploratory data analysis and principle component analysis visualisation was carried out using the DESeq2 package (Love, et al., 2014). Heatmaps were generated using the heatmap3 library (Zhao, et al., 2014), using the Spearman correlation coefficient between counts per million per gene as the distance metric for clustering.

3 replicates for each time point/tissue was run for RNA seq. Recent guidelines for “A survey of best practices for RNA-seq data analysis” indicates that “three replicates are the minimum required for inferential analysis” (Conesa, et al., 2016). We used the RNASeqPower library (Hart, et al., 2013) to perform power calculations on the count tables obtained after filtering out lowly expressed genes. The lowest median depth was observed in the E13.5S3 dataset, with a median number of 90 reads per tested gene in the library. Using this value as the coverage parameter for RNASeqPower, we revealed that with a within-group biological coefficient of variation of 0.1, which is commonly used for inbred animals, and with the 0.05 size of the test statistic we used (alpha) and a power of 0.95, we would have needed 1.14 samples per group to accurately quantitate differential gene expression with a logFC > 2. Hence, our analysis was adequately powered.

All tests used are in accordance with current best practices as outlined by Conesa et al. 2016 (Conesa, et al., 2016) and analysis are carried out in line with current best practices. All quality control metrics (% mapping, reads to genes etc) were consistent between the replicate samples.

### Gene ontology and pathway analysis

Gene ontology analysis was carried out using the topGO R package (Alexa, 2016) using the Fisher-elim algorithm (Alexa, et al., 2006). The Ensembl version 83 annotation was downloaded from biomaRt and used to generate the gene:category mappings. To take into account length bias in RNA-Seq gene ontology enrichment analysis (Young, et al., 2010) a list of genes expressed at similar levels but not differentially expressed between the two conditions was used, selecting 10 non-differentially expressed genes for every differentially expressed one. The union of these genes for each condition was used as the background list for topGO. For visualization, the results of the topGO analysis, expression values and the gene ontology mappings were exported to Cytoscape (Cline, et al., 2007), and the EnrichmentMap plugin (Merico, et al., 2010) was used for visualizing the enriched categories and the overlap between them. The size of each circle in Figure 1e and 1f indicates the number of genes contributing to each gene ontology category, while the thickness of the connections between circles indicates the degree of overlap between the two categories presented.

Ingenuity pathway analysis was used to identify upstream regulatory molecules such as transcription factors and miRNA. Genes that displayed an absolute logFC of >= 4 and an adjusted p-value of <= 0.01 were used as input, and only experimentally validated interactions described in animals, tissues and neuroblastoma cell lines were considered in the analysis. For the visualisations, the overlapping network of significant upstream regulators with over 10 regulated genes in the dataset was combined.

### Novel transcript prediction, annotation and differential expression testing

Sequencing reads were mapped using HISAT2 version 2.0.5 (Kim, et al., 2015) to the mm10 mouse genome, using gencode M8 to build an exon and splice junction reference. Stringtie 1.3.1c (Pertea, et al., 2015) was then used to assemble novel transcripts from these mapped reads. The minimum isoform abundance of the predicted transcript as a fraction of the most abundant transcript assembled at a given locus was required to be 0.05; transcripts were assembled with a length of 100 nt or greater; junctions with fewer than four bases on both sides were filtered out; and reads mapping closer than this 25 nt were merged together in the same processing bundle. FEELnc was then used to filter, predict the coding potential and annotate the assembled transcripts. First, monoexonic transcripts, transcripts that were less than 200 nt long, and biexonic transcripts in which one of the exons was less than 10 nt long were filtered out. Next, the random forest algorithm of FEELnc was trained on gencode M8 “protein_coding” transcripts as a set of coding genes, and gencode M8 “lincRNA”, “Mt_rRNA”, “Mt_tRNA”, “miRNA”, “misc_RNA”, “rRNA”, “scRNA”, “snRNA”, “snoRNA”, “ribozyme”, “sRNA”, and “scaRNA” as non-coding transcripts. K-mers of lengths 1,2,3,6,9 and ORF type 3 (containing either a start or a stop codon, or both) were used in training the model. A cutoff of 0.5268 was identified to be discriminatory for non-coding *vs*. coding transcripts, revealing 3666 novel putative non-coding and 1970 novel putative coding transcripts, originating from 4291 loci in the genome (Figure S7b). Finally, transcripts were classified using the FEELnc classifier with a window of 10000 nt proximity, using all transcripts in Gencode M8 for classification.

Gene level count tables were then generated for these transcripts (combining the gencode annotation and novel transcript coordinates) using featureCounts, and differential expression carried out as described above for the annotated transcripts, with both novel and annotated transcripts used for normalisation and differential expression testing.

### Single cell RNA sequencing

Cell suspension in a concentration of 600-1000 cells/*μ*L was used. C1 Suspension Reagent was added (all ‘C1’ reagents were from Fluidigm, Inc.) in a ratio of 4*μ*L to every 7*μ*L cell suspension. 11*μ*L of the cell suspension mix was loaded on a C1 Single-Cell AutoPrep IFC microfluidic chip designed for 10- to 17*μ*m cells, and the chip was then processed on a Fluidigm C1 instrument using the ‘mRNA Seq: Cell Load (1772x/1773x)’ script (30 min at 4°C). The plate was then transferred to an automated microscope (Nikon TE2000E), and a bright-field image (20× magnification) was acquired for each capture site using *μ*Manager (http://micro-manager.org (2)), which took <15 minutes. Quality of cells and control for doublets and processing of C1 chips were processed as described in Zeisel et al 2015 (Zeisel, et al., 2015) and Marques et al 2016 (Marques, et al., 2016).

List of all the single cell experiments performed:

2 E13.5 Brain experiments: 2 and 4 Fluidigm chips, respectively

1 E13.5 Spinal cord experiment – 4 Fluidigm chips

1 P7 Brain experiment – 4 Fluidigm chips

2 P7 Spinal cord experiments – 1 and 4 Fluidigm chips, respectively

2 experiments with P7 /P8 brain from E13.5 derived cells – 6 Fluidigm chips

1 experiment with P7/P8 brain from P3 derived cells – 3 Fluidigm chips

1 experiment with E17.5 brain – 1 Fluidigm chip

To prevent batch effects, in a balanced study design cells from different timepoints and regions would be mixed in each processed single cell run. Since we were collecting several timepoints and regions and using the Fluidigm C1 96 well Chips, this was not possible to implement. Cells from the same area and age were collected and processed at several time points. We did not observe differences between the experiments that would dramatically affect the analysis of the data, as assessed by comparison of the original dataset with our dataset where the confounding factors were regressed out.

### Single cell clustering

#### Quality control

Prior to clustering, cells from our dataset and the OPC, COP, and VLMC cells from Marques. et. Al (Marques, et al., 2016) were selected based on a minimum transcript threshold. Cells had to express a minimum of 1000 mRNA molecules per cell excluding mitochondrial RNA (filtered by the string “Mt-”), and repeat RNA (filtered by the string “r_”) in order to be considered for analysis. Distributions of transcript counts and total gene counts were calculated and categorized by cluster (Figure S4a). Additionally 84 cells were considered doublets due to a joint expression of either OL and neuronal genes, or a joint expression of OPC and VLMC associated genes. Weak single cell data / dead cells which did not pass the quality control check have been filtered away from the single cell analysis.

The number of cells used in single cell RNA-Seq to identify cell populations/states is constantly evolving. The number of cells used in this study (1974 cells post-quality control) is considerable higher than recent published studies using similar technologies to investigate neural progenitor states (for examples, 272 cells in Telley et al, 2016 (Telley, et al., 2016)).

#### BackSPIN clustering

The post QC dataset was clustered using the BackSPIN2 algorithm as previously described (Marques, et al., 2016; Romanov, et al., 2016). In short, the algorithm is an adaptation to the sorting algorithm SPIN (Tsafrir, et al., 2005) wherein a bi-clustering is performed by sorting the cells and genes into a one-dimensional ordering where a binary split is performed based on the distribution of genes within each ordering. The algorithm repeatedly performs feature selection and subsequent splits until a certain threshold is achieved.

#### Pathway and geneset overdispersion analysis (PAGODA)

The post QC dataset was clustered using PAGODA (SCDE R-package) (Fan, et al., 2016). First, the drop-out rate is determined and the amplification noise is estimated through the use of a mixture-model. Then the first principal component is calculated for all gene sets and GO-term clusters are provided or identified, and over-dispersion is defined as the amount of variance explained by the gene set above expectation. Holm procedure is used as a part of the SCDE package.

#### Unbiased cluster determination PAGODA

Cluster generation within PAGODA is based on hierarchical clustering of the distance measure obtained from the first principal component of the gene sets. We developed an algorithm to determine the final cluster amount in an unbiased way. First, we perform a differential expression analysis using SDCE for each new split to calculate for each cluster what genes are differentially expressed between them, we filter genes by a requirement to be expressed in at least 60% of the population, and then we calculate a p-value assuming a normal distribution based on the Z-scores obtained from the differential expression analysis. Subsequently we either discard the split or accept the split based on the significance of the top 20 most significant genes (p<0.01). Using these settings the algorithm returned 15 clusters from the dataset.

#### Merging of BackSPIN2 and PAGODA clusters

In general, PAGODA and BackSPIN2 gave rise to clusters with similar transcriptional profiles. While we adopted the main BackSPIN2 clusters, PAGODA resolved OPCs as three populations and not as one population (as in BackSPIN2). This might be due to low expression and the possible quiescent phenotype of some of the OPC clusters. PAGODA has the advantage of looking at patterns of groups of genes instead of individual genes thus increasing sensitivity for low expressing cells. Therefore we retained the three found OPC clusters in PAGODA and the NP3 cluster from PAGODA (due to their low expression) and merged them with the remaining BackSPIN2 clusters. The remaining cells classified by backSPIN2 as OPCs but not by PAGODA, were reassigned to the COP and NP3 clusters, given their transcriptional profile.

BackSPIN2 clustering indicated distinct clusters within the NFOL cluster. Enrichment analysis and marker selection revealed shared markers and enriched genes as well as some minor differences in genes such as *Mog* and *Mag* expression. Further analysis of these subclusters did not fall within the scope of this paper and as such the clusters were merged into the NFOL cluster. Furthermore, Enrichment analysis and marker selection of the NP1 cluster revealed a clear bimodal expression profile, hence we used the BackSPIN2 clustering data to split one level deeper within the NP1 population resulting in the NP1a and NP1b subclusters. PAGODA also revealed these when allowed to oversplit, our splitting algorithm indicated this split to form valid clusters. However, the NP1 cluster split was preceded by a number of non-valid splits, meaning that this cluster is a subcluster and statistical evaluation of split validity is less effective due to the small cluster size.

### Single cell RNA sequencing pathway analysis

Single cell differential gene expression results were obtained using SCDE, and a p-value was calculated assuming a normal distribution based on the Z-scores obtained. Genes were filtered by the requirement that they should be expressed in at least 20% of the population to avoid highly significant results driven by expression in a few cells within the population. The uncorrected Z-score and p-value were then used to filter the list of differentially expressed genes in Ingenuity pathway analysis, requiring the abs(Z-score) to be > 4 and the p-value to be <= 0.01. Similarly to the analysis of the bulk sequencing data, only experimentally observed interactions described in animals, tissues and neuroblastoma cell lines were considered in the analysis. Full results are presented in Table S6.

### Single-cell near-neighbour network embedding (SCN3E)

#### Feature selection

Initial gene selection involves a cutoff for genes expressing at least 1 transcript, then a feature selection is performed using the coefficient of variation, and for each gene we select genes above the support vector regression fitted line.

#### Binarization of expression data

In order to reduce noise and to define epigenetic states a binarization is performed in where the expression matrix is reduced to a binary representation where a value of 1 indicates that the gene is on and a value of 0 indicates that the gene is off. This binary cell barcode is used for subsequent correlation analysis reducing the influence of differences caused by total mRNA content fluctuations in cells of otherwise similar states.

#### Dimensional reduction of both binary and count based expression matrix spaces

For dimensional reduction we use diffusion mapping, a non-linear dimensional reduction technique (Rpackage DPT) (Haghverdi, et al., 2016) including a locally scaled transition matrix for improved resolution. The first 7 diffusion components are used for subsequent network embedding (depending on the complexity of the dataset). For correlation we use the first 6 diffusion components of the reduced binary expression space, this is a relatively low number of dimensions which was chosen to reduce any noise in the correlation and also because the binary expression space is inherently less complex compared to the full expression space.

#### Nearest neighbor network embedding

Distances between cells are defined in Euclidean space based on the previously defined diffusion map coordinates from the gene expression space. Per individual cell, the 20 nearest neighbors are calculated and network edges are created between them. Edges are then weighted based on the Pearson correlation as defined by the 6 diffusion components of the binary gene space. The edge weights are then raised to a high power (we used 10 preventing any cells from becoming detached from the network) and edge weights of less than 0.1 are removed, leaving only strongly correlating edges without losing cells from the network.

### Single cell weighted correlation network analysis (WGCNA)

A signed gene co-expression network was constructed using the WGCNA package in R (Langfelder and Horvath, 2008). The standard workflow was followed as described in the WGCNA tutorial
(https://labs.genetics.ucla.edu/horvath/CoexpressionNetwork/Rpackages/WGCNA). Genes with less than 5 counts for any cell were discarded from the analysis. Soft thresholding estimation was performed and a gene module size of 20 genes was defined for the clustering of genes by topological overlap distance. Node centrality was used to determine the top putative regulators for each module (Table S6). Correlation is performed between each individual cell and then averaged over each cluster. The minimum group size is 14 cells hence justifying averaging. WGCNA sample size is 15 samples, thus 1974 cells is significantly above the number recommended for the workflow.

### Lineage tracing

Pdgfra-cre-ERT/RCE (Kang, et al., 2010) (mixed C57BL/6NJ and CD1 background) female mice were injected 2mg tamoxifen (20mg/ml, Sigma) at pregnancy day E.12.5 (txE12.5) or when pups were P3 (txP3). Brain tissue was then harvested at P7-P8 for electrophysiology and single cell RNA sequencing and at P21 for immunohistochemistry experiments, respectively.

No further behavioral experiments were done in this group of animals.

### Electrophysiology

P7-P8 pups were deeply anesthetized with isoflurane and brains were collected in ice-cold solution of the following composition (in mM): 62.5 NaCl, 100 sucrose, 2.5 KCl, 25 NaHCO_3_, 1.25 NaH_2_PO_4_, 7 MgCl_2_, 1 CaCl_2_, and 10 glucose. Brains were vibratome sectioned to 300μm slices in the same solution and were then let recover for 1 h at room temperature in oxygenated aCSF (in mM): 125 NaCl, 2.5 KCl, 25 NaHCO_3_, 1.25 NaH_2_PO_4_, 1 MgCl_2_, 1 CaCl_2_, and 10 glucose. To maximize the selection of OPCs for electrophysiological recordings, we targeted small circular eGFP+ cells found in the corpus callosum, especially avoiding elongated cells attached to blood vessels (which would be VLMCs or pericyte-lineage cells, labeled in this mouse line).

Whole-cell patch-clamp recordings were performed at 25±2°C, with slices continuously perfused with oxygenated aCSF. Patch electrodes were made from borosilicate glass (resistance 5–8 MΩ; Hilgenberg, GmbH) and filled with a solution containing (in mM): 130 CsCl, 4 NaCl, 0.5 CaCl_2_, 10 HEPES, 10 EGTA, 4 MgATP, 0.5 Na_2_GTP. Neurobiotin (0.5%, Vectorlabs) was included for post-hoc identification of recorded cells.

Cells were recorded in voltage-clamp mode held at -70 mV. At the end of the recording tetrodoxin (TTX, 10μM) or CNQX/MK-801 (10μM /5μM) were applied for 10 min to block spontaneous neuronal activity or glutamatergic inputs respectively. Currents were recorded with an Axopatch 200B amplifier (Molecular Devices), sampled at 10 kHz and digitized with Digidata 1322A (Molecular Devices). All drugs were ordered from Sigma. In order to confirm their OPC identity, slices were fixed with 4% PFA for 1-2 h after recording, washed and kept in PBS 4°C until stained for NG2 (rabbit anti-NG2 1:200, Millipore), Streptavidin 555 (1:1000, Invitrogen) and Alexa-647 anti-rabbit (1:400, ThermoFisher). We did not recover post-staining for every recorded cell included in the analysis. Nevertheless, we enriched our sample for OPCs, using a reporter mouse line and targeting small round eGFP+ cells. Accordingly, all cells have homogenous intrinsic properties (e.g., input resistance – which differs greatly in more mature stages). Furthermore, in relation to synaptic inputs we recovered eGFP+/NG2+ cells comprising both the lowest and the highest frequencies of synaptic events.

7 slices from 2 txE13.5 animals and 15 slices from 4 txP3 animals were recorded. In electrophysiology it is common to recorder from 6-10 cells per group. After a first batch of experiments, we appreciated that the txP3 group was more heterogeneous (number of events) and we decided to enlarge the sample to 15 cells in order to confirm the synaptic events distribution.

### Electrophysiology analysis

All traces were low-pass filtered at 1KHz (8-pole Bessel filter) and only events with amplitude larger than -10pA were included in the analysis. We utilized a semi-automated event-detection on Clampex with which events were visually inspected and unclear cases were discarded. Cells that died during recording or whose signals were not analyzed were excluded. For each cell, average amplitude and frequency values were calculated by total number events per time. Data are mean ± s.e.m. P-values are from Student’s two independent sample (p=0.27; frequency F=13.95, t= -1.53, df= 20; amplitude (F=0.99, t= 1.1, df = 18)) or paired t-tests (p=0.033, p=0.06 and p=0.11; paired txE12.5 x txE12.5+TTX (t=2.497, df=4); paired txP3 x txP3+TTX (t=2.4974, dff=2); paired txP3 x txP3+CM (t=3.71, dff=3)).

### Immunohistochemistry

Pdgfra-CreERT/RCE mice recombined with Tamoxifen at E12.5 were perfused at P20 with PBS followed by 4% PFA. Brains and spinal cords were dissected and postfixed with 4% PFA for 2h, at 4°C. The tissues were then cryoprotected with a 30% sucrose solution for 48 hours. The tissues were embedded into OCT (Tissue-Tek) and sectioned coronally (20 um thickness).

Sections were quickly boiled in antigen retrieval (Dako, S1699) and stood in the antigen solution until cooling down. They were then permeabilized in PBS/0.3% Triton (Millipore) 3× 5 minutes and blocked for 1 hour in PBS/0.3% Triton/5% normal donkey serum (Sigma, D9663) at room temperature. Sections were incubated overnight at 4°C with primary antibodies [GFP (Abcam, ab13970, chicken 1:2000), PDGFRA (R&D, AF1062, Goat 1:200), COL1A1 (Abcam, ab21286, rabbit, 1:50) or CC1 (anti-APC; Millipore, OP80, Mouse 1:100) diluted in PBS/0.3% Triton/2% normal donkey serum. After washing the sections 3x 5min with PBS, secondary Alexa Fluor-conjugated antibodies (Invitrogen, Alexa Fluor 488 1:500, Alexa Fluor 555 1:1000 and Alexa Fluor 647 1:250) diluted in PBS/0.3% Triton/2% normal donkey serum were added and incubated for 1 hour at room temperature. Thereafter, slides were mounted with mounting medium containing DAPI (Vector, H-1200) and kept at 4°C until further microscopic analysis.

Antibodies have been cited by other authors. Lists are available on the webpage of the provider company or have been tested for immunohistochemistry in mouse by the company.

### Microscopy and analysis

Combined images of DAPI, Alexa 555, Alexa 488 and Alexa 647, spanning the corpus callosum (CC) and dorsal horn were obtained in a Zeiss LSM700 Confocal. For quantifications, 3 animals were used in each timepoint and 4-5 slices were photographed per animal. An average of 33 and 46 photos in CC and dorsal horn respectively were counted per animal. The number of animals used was similar to those reported in previous publications presenting similar experiments (Kessaris, et al., 2006). All the countings were normalized to the area analysed in each photo. % of Pdgfra+Col1a1- (OPCs) and Pdgfra+Col1a1+ (VLMCs) cells out of GFP were calculated. The remaining GFP cells (Pdgfra-/Col1a1-) were considered as non-OPC cells. The percentage of GFP cells out of the Pdgfra+ population was counted as being OPCs derived from the E13.5 wave while the remaining Pdgfra+GFP- were considered to derive from the second/third waves.

### Cell culture

OLI-neu, an immortalized oligodendrocyte precursor cell line (Jung, et al., 1995), was cultured in DMEM (Gibco, Germany) 1x Penicillin/Streptavidin (Gibco), 0.1% N2 supplement (Gibco), 400ng/mL T4, 340ng/mL T3 (Sigma), lOng/mL bFGF and lOng/mL PDGF-BB (Peprotech). Cells were grown in flasks or plates coated on poly-L-lysine (PLL; Sigma) for at least 30min. This cell line is not in the ICLAC or NCBI biosample databases and tested negative for mycoplasma.

### Knockdown of ncRNA

Cells were seeded in proliferative media the day prior to experiment in 6 multiwell plates coated with PLL, at a density of 250 000 cells per well. A mix of 3μg (equivalent to lOOnM) of custom siRNA (Dharmacon) was added to the cells in a complex with Lipofectamine 2000 for 4h in Optimem (ThermoFisher). Medium was then changed to proliferative medium and 20h later cells were collected in Qiazol (Qiagen) for RNA extraction.

siRNA sequence:

Sense-GUACACAGGGAAAGGUCUAUU

Antisense- UAGACCUUUCCCUGUGUACUU

### Real time quantitative PCR analysis

Total RNA was extracted using miRNeasy micro kit from Qiagen. 500ng of RNA was converted to cDNA using High-Capacity cDNA Reverse Transcription Kit with RNase Inhibitor (Applied Biosystems). Real time qPCR analysis was performed with Fast sybr green mastermix (Applied biosystems) in the 7900HT Fast System equipment. Tbp, Ubc and Gapdh were used as housekeeping genes and the geometrical mean was calculated for normalization of the relative expression values.

Primers used for each gene are as follows:

Tbp F- GGGGAGCTGTGATGTGAAGT;

Tbp R- CCAGGAAATAATTCTGGCTCA

Ubc F- AGGTCAAACAGGAAGACAGACGTA

Ubc R- TCACACCCAAGAACAAGCACA

Gapdh F- GAG AAA CCT GCC AAG TA

Gapdh R- AGA CAA CCT GGT CCT CA

Pcdh17IT F- GCA ATC ACT AAC AGA ATT CCG TGT

Pcdh17IT R- TTG TAA ATA GCA GTA GGG AAG GG

Pcdh17 F- AAG ACA CTA ACA AAG GCT CCT GCT

Pcdh17 R- CTA TCA GAA TGA CCA AGC ACT CG

SoxlO F- AGATGGGAACCCAGAGCAC

SoxlO R- CTCTGTCTTTGGGGTGGTTG

Mog F- ATGAAGGAGGCTACACCTGC

Mog R- CAAGTGCGATGAGAGTCAGC

Myrf F- GGA GGC TTT TTG CAG AGG TT

Myrf R- AAT GTC ACC CGG AAG TGG TA

Egr1 F- AGC GCC TTC AAT CCT CAA G

Egr1 R- TTT GGC TGG GAT AAC TCG TC

Tcf712 F- CCT GAC GAG CCC CTA CCT

Tcf712 R- TGC AAT CTG GTG TTC AAT CG

### smFISH method

smFISH was carried out as previously described (Zeisel, et al., 2015). The cells were permeabilized using PBS-TritonX 0.5% for 10 min at room temperature followed by 24 hours of hybridization with 250 nM fluorescent label probes (Biosearchtech, Petaluma, CA, USA) at 38.5°C and counterstained with DAPI (Life Technologies). The sections were mounted with pro-long gold (Life Technologies) and image stacks (0.3 μm distance) were acquired using a Nikon Ti-E with motorized stage (Nikon). The images were analyzed with a custom python script using the numpy, scipy.ndimage (Jones E, Oliphant E, Peterson P, et al. SciPy: Open Source Scientific Tools for Python, 2001) and scikit-image) libraries. Briefly, after background removal using a large kernel gaussian filter, a Laplacian-of-Gaussian was used to enhance the RNA dots. This experiment was done once to confirm the qPCR results indicating successful knockdown.

## Acknowledgements

We would like to thank Amit Zeisel, Alessandra Nanni, Ahmad Moshref and Johnny Söderlund, and the Single Cell Genomics Facility at SciLifeLab for support. DV would like to acknowledge the University of Sydney HPC service at The University of Sydney for providing HPC resources that have contributed to the research reported in this paper. This work was supported in part by a University of Sydney HPC Grand Challenge Award. Access to IPA was provided by the University of Sydney’s Mass Spectrometry Core Facility. DV was supported in part by a Boehringer Ingelheim Travel Grant. Work in RJT lab was supported by Australian NHMRC grant APP1043023. LV, SG and ÅB are supported by a grant from the Knut and Alice Wallenberg Foundation to the Wallenberg Advanced Bioinformatics Infrastructure. Work in SL lab is supported by Knut and Alice Wallenberg Foundation grant 2015.0041 and Swedish Foundation for Strategic Research (RIF14- 0057). J.H-L is supported by VR, StratNeuro, Hjärnfonden, and EU FP7/Marie Curie Actions. A.M.F. by the European Committee for Treatment and Research of Multiple Sclerosis (ECTRIMS). Work in G.C.-B.’s research group is supported by Swedish Research Council, European Union (FP7/Marie Curie Integration Grant EPIOPC, Horizon 2020 European Research Council Consolidator Grant EPIScOPE), Swedish Brain Foundation, Ming Wai Lau Centre for Reparative Medicine, Swedish Cancer Society (Cancerfonden), Petrus och Augusta Hedlunds Foundation and Karolinska Institutet. R.J.T. is an employee of Illumina Inc.

